# Recurrent duplication and diversification of a vital DNA repair gene family across Drosophila

**DOI:** 10.1101/2023.10.04.560779

**Authors:** Cara L. Brand, Genevieve T. Oliver, Isabella Z. Farkas, Mia T. Levine

## Abstract

Maintaining genome integrity is vital for organismal survival and reproduction. Essential, broadly conserved DNA repair pathways actively preserve genome integrity. However, many DNA repair proteins evolve adaptively. Ecological forces like UV exposure are classically cited as drivers of DNA repair evolution. Intrinsic forces like repetitive DNA, which can also imperil genome integrity, have received less attention. We recently reported that a *Drosophila melanogaster*-specific DNA satellite array triggered species-specific, adaptive evolution of a DNA repair protein called Spartan/MH. The Spartan family of proteases cleave hazardous, covalent crosslinks that form between DNA and proteins (“DNA-protein crosslink repair”). Appreciating that DNA satellites are both ubiquitous and universally fast-evolving, we hypothesized that satellite DNA turnover spurs evolution of DNA-protein crosslink repair beyond *D. melanogaster*. This hypothesis predicts pervasive Spartan gene family diversification across the Drosophila phylogeny. To study the evolutionary history of the Drosophila Spartan gene family, we conducted population genetic, molecular evolution, phylogenomic, and tissue-specific expression analyses. We uncovered widespread signals of positive selection across multiple Spartan family genes and across multiple evolutionary timescales. We also detected recurrent Spartan family gene duplication, divergence, and gene loss. Finally, we found that ovary-enriched parent genes consistently birthed testis-enriched daughter genes. To account for Drosophila-wide, Spartan family diversification, we introduce a mechanistic model of antagonistic coevolution that links DNA satellite evolution and adaptive regulation of Spartan protease activity. This framework, combined with a recent explosion of genome assemblies that encompass repeat-rich genomic regions, promises to accelerate our understanding of how DNA repeats drive recurrent evolutionary innovation to preserve genome integrity.

## INTRODUCTION

Our genome is under constant threat from both intrinsic and extrinsic DNA damaging agents. Cellular metabolites, DNA replication errors, radiation, and chemical mutagens, among others, trigger DNA lesions. A suite of pathways dedicated to repairing DNA damage actively preserve genome integrity. These vital pathways are found in virtually all eukaryotes, from humans to worms to yeast (Taylor and Lehmann 1998). Despite the essentiality and deep conservation of eukaryotic DNA repair, many DNA repair proteins are unconserved. These repair proteins evolve rapidly under positive selection, resulting in highly divergent alleles between closely related species (Sawyer and Malik 2006; Demogines, et al. 2010; Lou, et al. 2014; Lee, et al. 2016; Lou, et al. 2016; Sun, et al. 2018; Zhang, et al. 2019; Kolora, et al. 2021; Rowley, et al. 2021; Zhang, et al. 2023). This rapid evolution has been attributed largely to extreme environments and viral pathogens; however, a building literature implicates rapidly evolving DNA repeats as dynamic substrates that select for innovation of DNA repair proteins (Brand and Levine 2021).

Genomic regions rich in DNA repeats are dominated by fast-evolving transposons and fast-evolving, tandemly repeating units called satellite DNA. Transposons and satellites frequently engage DNA repair pathways. Transposons hijack host DNA repair machinery to complete integration into a novel genomic location (Feschotte and Pritham 2007). Consequently, mutations in key DNA repair genes either increase or decrease transposon activity, consistent with DNA repair-mediated transposon restriction (Downs and Jackson 1999; Scholes, et al. 2001; Griffith, et al. 2003; Gasior, et al. 2008; Suzuki, et al. 2009; Mita, et al. 2020). DNA breaks in satellite arrays are repaired by a specialized DNA damage response pathway that prevents catastrophic recombination between non-homologous genomic regions (Chiolo, et al. 2011; Ryu, et al. 2015; Feng, et al. 2016). This engagement of fast-evolving transposons and fast-evolving satellites with DNA repair pathways raises the possibility that preserving repetitive DNA integrity requires recurrent evolutionary innovation of DNA repair factors (Brand and Levine 2021).

To date, the only empirical evidence that DNA repeats trigger adaptation at DNA repair machinery comes from the DNA-protein crosslink repair pathway (Brand and Levine 2022). Transient and reversible crosslinks form between DNA and the proteins that perform various DNA-templated functions (Figure 1A). However, mutagens and enzymatic errors can render these reversible crosslinks irreversible (Stingele, et al. 2017). Irreversible DNA-protein crosslinks block essential DNA transactions such as replication and transcription (Fu, et al. 2011; Nakano, et al. 2012). Specialized DNA repair proteins, called Spartan proteases, resolve these harmful lesions by degrading the crosslinked protein (Lopez-Mosqueda, et al. 2016; Stingele, et al. 2016; Vaz, et al. 2016)). This protease activity is highly regulated: overactive DNA-protein crosslink repair disrupts vital DNA-protein interactions while insufficient repair blocks essential DNA transactions (Stingele, et al. 2016). Intriguingly, Spartan protease regulation appears to coevolve with DNA repeats. A recent, evolution-guided functional analysis revealed that the *Drosophila melanogaster* Spartan homolog, called Maternal Haploid (MH), antagonistically coevolved with a DNA satellite that proliferated along the *D. melanogaster* lineage (Brand and Levine 2022). Replacing the native *D. melanogaster* gene with an adaptively diverged *mh* allele induces extensive DNA damage, ostensibly caused by over-active DNA-protein crosslink repair. Deletion of the *D. melanogaster*-specific DNA satellite array inhibited DNA damage. These data suggest that DNA satellite proliferation perturbed healthy DNA-protein crosslink repair and that MH adaptive evolution mitigated this perturbation.

**Figure 1.**
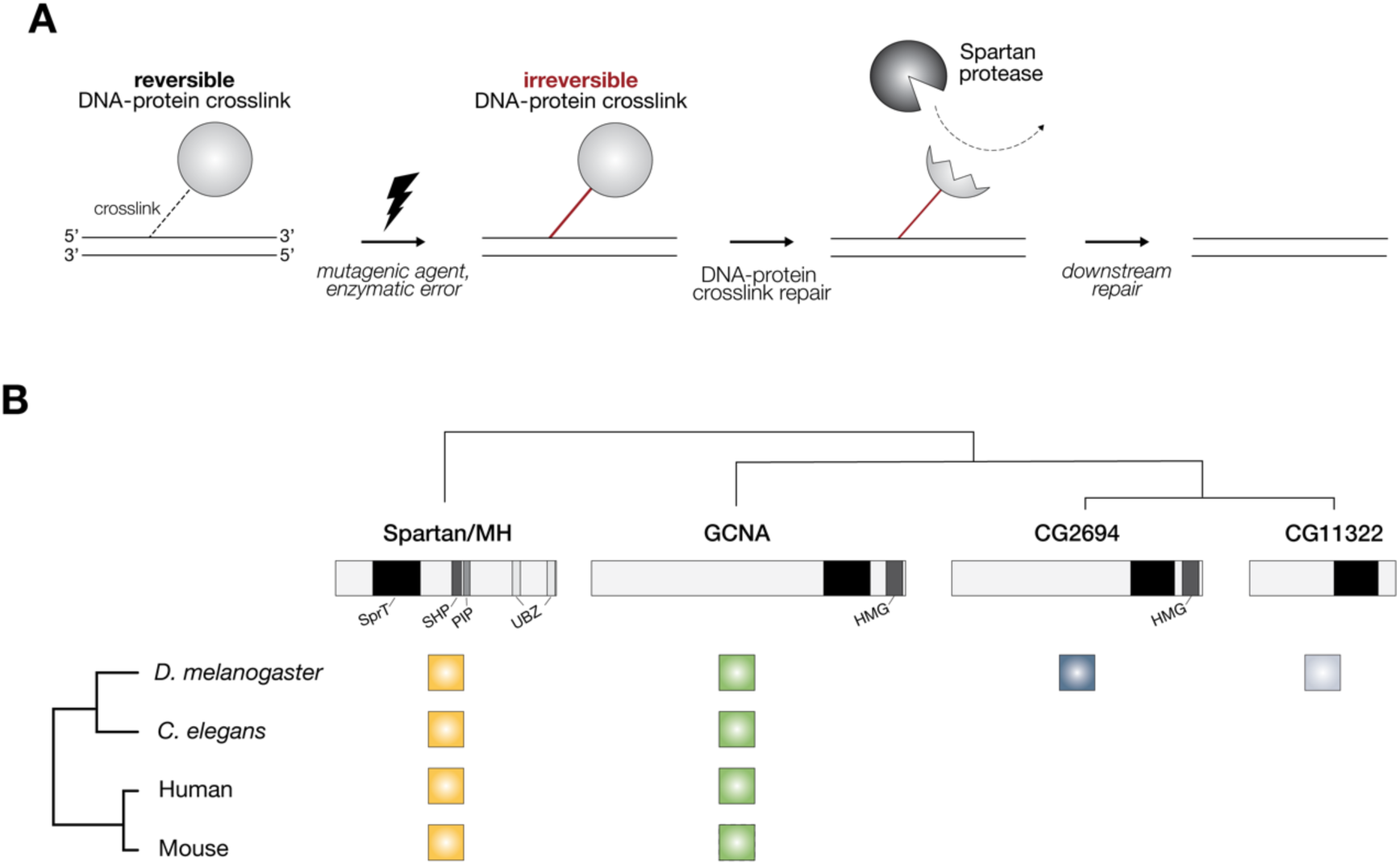
Spartan family proteases are ancient and repair DNA-protein crosslinks. (**A**) Transient and reversible crosslinks form between DNA and proteins that perform a variety of genome functions. Mutagens and enzymatic errors can render these reversible DNA-protein crosslinks irreversible. Irreversible crosslinks trap proteins on the DNA, blocking vital DNA-dependent functions. Spartan proteases cleave proteins trapped as DNA-protein crosslinks. The remaining structure is bypassed by translesional synthesis and further resolved by downstream DNA repair pathways (Larsen, et al. 2019). (**B**) Spartan family proteases are present in worms, insects, and mammals. The Spartan family is defined by the SprT protease domain. Spartan/MH homologs (yellow) encode additional defined domains that regulate protease activity. GCNA homologs (green) encode an N-terminal intrinsically disordered region and a non-canonical HMG box (Carmell, et al. 2016). Two additional GCNA-like proteases, CG2694 (blue) and CG11322 (light blue), are restricted to Drosophila.

Motivated by this specific case of antagonistic coevolution between a single Spartan protease and a single DNA satellite, we hypothesized that satellite evolution more generally perturbs DNA-protein crosslink repair. Satellites are ubiquitous across Drosophila (and all eukaryotes, (Britten and Kohne 1968)) and DNA satellite-rich genomic regions consistently represent some of the most diverged sequence between closely related species (Iwata-Otsubo, et al. 2017; Jagannathan, et al. 2017; Wei, et al. 2018; Cechova, et al. 2019; de Lima, et al. 2020; Sproul, et al. 2020; de Lima and Ruiz-Ruano 2022). The pervasiveness and rapid evolution of satellite DNA predicts diversification of DNA-protein crosslink repair machinery across the Drosophila phylogeny – not only of *mh* but also of other Spartan gene family members (Fielden, et al. 2018; Bhargava, et al. 2020).

To comprehensively explore Spartan family diversification, we conducted population genetic, molecular evolution, phylogenomic, and expression analyses of the Drosophila Spartan family. *D. melanogaster* encodes four Spartan proteases defined by a shared “Spartan-like protease” domain with a conserved catalytic core: *Germ cell nuclear acidic peptidase* (*Gcna*) and *mh* are characterized and shared across eukaryotes (Carmell, et al. 2016; Fielden, et al. 2018) while *CG2694* and *CG11322* are restricted to Drosophila and are functionally uncharacterized (Bhargava, et al. 2020); Figure 1B). All four genes are enriched in either the female or male germline tissue (Dai, et al. 2006; Delabaere, et al. 2014; Bhargava, et al. 2020). Here we report signatures of positive selection at all four Spartan family proteases in addition to dynamic whole gene births and deaths across the Drosophila phylogeny. We discovered that Spartan gene paralogs also undergo gene expression evolution, with repeated evolutionary transitions from ovary-to testis-biased expression across parent and daughter genes. These data support our prediction of Spartan family-wide, and Drosophila genus-wide, adaptive diversification. We propose a model under which DNA satellite expansions and contractions alter the abundance of single-/double-stranded DNA junctions (“ss/dsDNA junctions”), a DNA structure that stimulates Spartan protease activity (Reinking, et al. 2020). Changes to the abundance of ss/dsDNA junctions perturb protease activity, and ultimately, healthy rates of DNA-protein crosslink repair. Adaptive Spartan protease evolution mitigates the deleterious consequences of either too much or too little DNA-protein crosslink repair. This model of coevolution between DNA satellites and Spartan proteases offers a mechanistic understanding for how an ever-changing landscape of DNA satellites selects for DNA repair diversification.

## RESULTS AND DISCUSSION

### Recent and recurrent evolution of Spartan family genes

To detect evidence of recent adaptive sequence evolution, we performed a molecular population genetic analysis of *D. melanogaster* and *D. simulans*, sisters species that diverged ∼2.5 million years ago (Garrigan, et al. 2012). Specifically, we compared synonymous and nonsynonymous polymorphisms, within *D. melanogaster* and within *D. simulans* populations, to synonymous and nonsynonymous divergence between the two species. Under the McDonald-Kreitman test framework (McDonald and Kreitman 1991), an excess of nonsynonymous divergence is consistent with a history of adaptive evolution. We detected adaptive evolution between *D. melanogaster* and *D. simulans* at three of the four Spartan family genes (Table 1). Furthermore, polarized McDonald-Kreitman tests revealed evidence of adaptive evolution along one or both species’ lineages (Table S1).

**Table 1:**
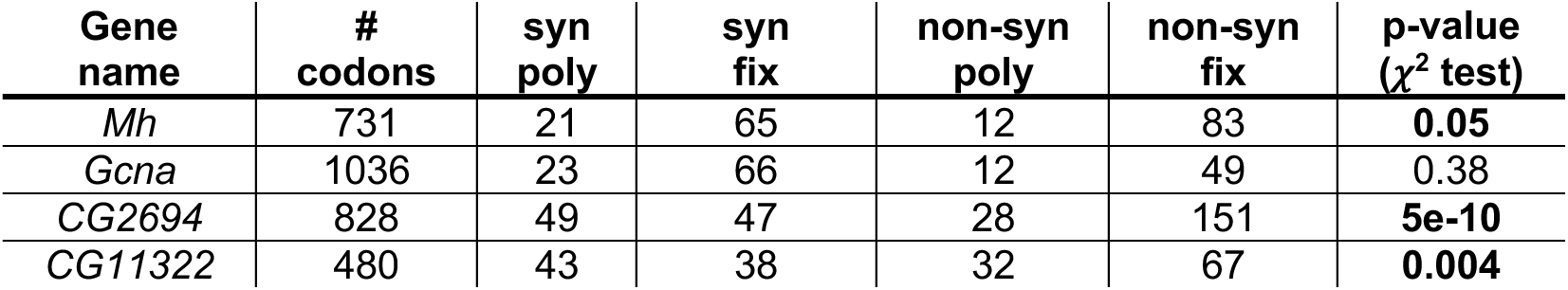
Multiple Spartan family genes evolve adaptively on recent evolutionary timescales. Counts of synonymous (syn) and nonsynonymous (non-syn) polymorphic (poly) and fixed (fix) sites within and between *D. melanogaster* and *D. simulans*.

To comprehensively explore the inference that adaptive evolution of Spartan family genes occurs along multiple species lineages, we investigated rates of molecular evolution over longer timescales. To limit synonymous site saturation across the Drosophila phylogeny (Larracuente, et al. 2008; Kumar, et al. 2017) while capturing deeply divergent lineages, we analyzed three distinct *Drosophila* clades in parallel: the *melanogaster* group (*n* = 23, ∼25 million years diverged), the *obscura* group (*n* = 12, ∼12 million years diverged), and the *virilis-repleta group* (*n* = 5, ∼25 million years diverged; Figure 2). Using a phylogeny-based, maximum likelihood approach to investigate molecular evolution ((Yang 1997); see Methods), we identified Spartan family genes with elevated rates of nonsynonymous substitutions relative to synonymous substitutions (*dN/dS*). We note that the restriction of *CG11322* to only a few *melanogaster* group species disabled us from including this young gene duplicate in our analysis (Dai, et al. 2006).

**Figure 2.**
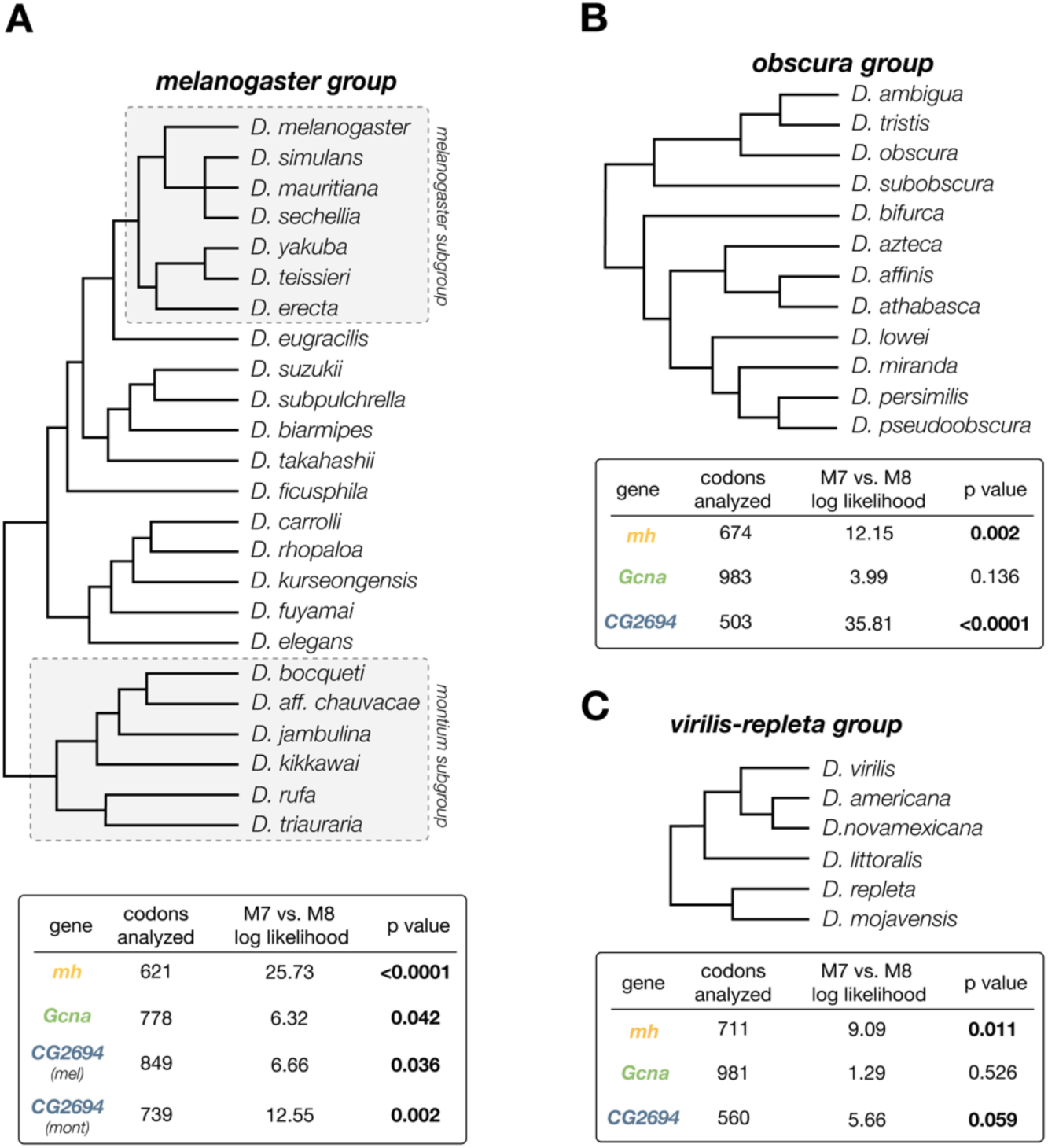
Spartan family proteases evolve under positive selection across multiple Drosophila clades. Phylogenetic relationship of species analyzed within the (**A**) *melanogaster*, *obscura*, and (**C**) *virilis-repleta* groups and the corresponding results from the molecular evolution analysis. Below each species tree are the results from an analysis of alternative NSsites models in the software package, PAML (see Methods). An unreliable alignment of *CG2694* orthologs across the *melanogaster* group required us to run separate molecular evolution analyses on the *melanogaster* and the *montium* subgroups.

Molecular evolution analysis of the Spartan gene family across three distinct clades revealed a heterogenous pattern of recurrent positive selection across genes and clades (Figure 2). We found that both *mh* and *CG2694* evolve adaptively in the *melanogaster* group, in the *obscura* group, and in the *virilis-repleta* group. In contrast to *mh* and *CG2694*, we detected weak evidence of positive selection at *Gcna* across to the *melanogaster* group but no evidence of positive selection across the *obscura* and the *virilis-repleta* groups. In summary, our analyses of adaptive sequence evolution across both short and long timescales, across multiple distinct clades, and across multiple members of the Spartan gene family, suggest that DNA-protein crosslink repair is under pressure to innovate by a pervasive and persistent evolutionary force.

### Recurrent birth of young Spartan family genes

Codon evolution is only one mechanism of evolutionary diversification. Gene turnover by recurrent duplication and/or pseudogenization can also diversify gene family repertoires. Previous reports found that *CG11322* recently duplicated from *CG2694* (Dai, et al. 2006) and that *mh* tandemly duplicated in *D. simulans* and *D. mauritiana* (Chakraborty, et al. 2021). This precedent of recent Spartan gene duplication motivated our investigation of gene turnover across a deeper evolutionary time span. We took advantage of 42 highly contiguous Drosophila genome assemblies (Table S2) that enabled us to confidently assign orthologs and paralogs. To define a set of Spartan family members across this 40-million-year stretch of evolutionary time, we used a combination of tBLASTN, phylogenetic tree building, and synteny analysis (Figure 3A).

**Figure 3.**
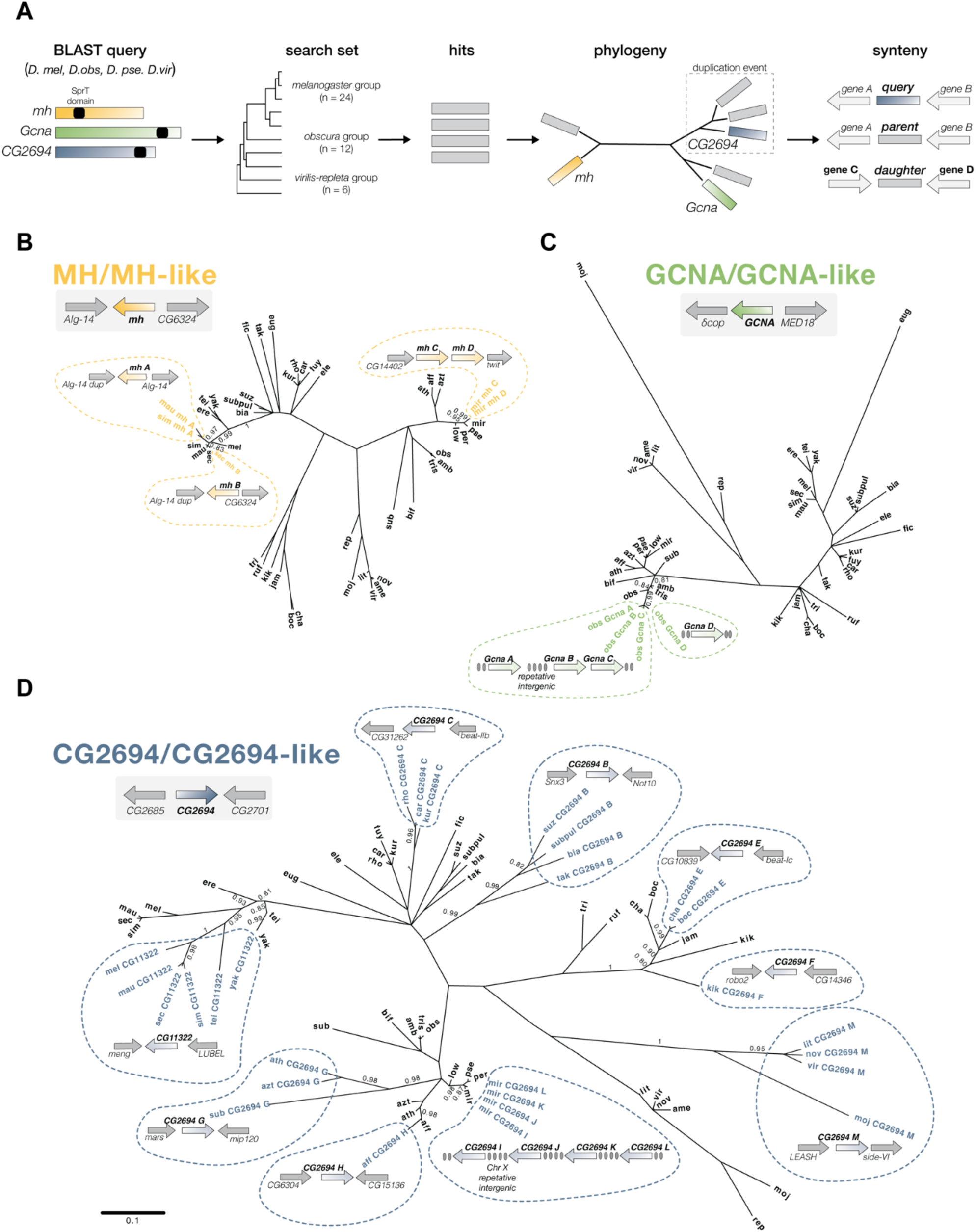
Heterogeneous duplication rates across the Spartan gene family. (**A**) Schematic of the pipeline used to identify additional Spartan proteases across Drosophila. We used the *D. melanogaster*, *D. obscura*, *D. pseudoobscura*, and *D. virilis* Spartan protease coding sequences as a query in a tBLASTn search of 42 highly contiguous Drosophila genome assemblies (Table S2). We built a phylogenetic tree of all significant BLAST hits using the SprT domain to infer Spartan gene subfamilies (Figure S2, Table S3). We then built phylogenetic trees of the (**B**) *mh Gcna* and (**D**) *CG2694* subfamilies (see Table S2 for full species names). In parallel, we subjected all hits to synteny analysis to delineate orthologs and paralogs within and between genome assemblies. Gray boxes show each Spartan family protease with the flanking genes delineated. Dashed bubbles show the lineage-restricted daughter genes with the flanking genes delineated. The four tandem *Gcna* duplicates in *D. obscura* are embedded in repetitive, intergenic sequences (grey ovals) that precluded precise mapping. The tandem *CG2694* duplicates in *D. miranda* are embedded in repetitive DNA that maps to a high-quality *X* chromosome assembly. Using synteny, we confirmed that the highly diverged *D. eugracilis* and *D. mojavensis Gcna* orthologs are represented on the phylogeny (Figure S3). Note that *CG2694 C* is not represented because it lacks a SprT domain (see Figure 4A). Only resolved nodes with branch support > 0.75 are shown. Branch support values indicated correspond to nodes of interest.

We identified 24 independent duplication events and, at minimum, 17 gene loss events. Like the adaptive sequence evolution reported above, these gene duplication events were heterogeneously distributed across *Gcna*, *mh*, and *CG2694* (Figure 3, 4). The *X*-linked *mh* duplicated independently at least five times but only along a restricted number of lineages. We detected three duplications in the *simulans* clade (including a previously reported duplication, (Chakraborty, et al. 2021); Figure 3B, 4A, S1), and two duplications along the lineage leading to *D. miranda* (Figure 3B, 4B). Similarly, *Gcna* duplicated four times along the branch leading to *D. obscura* (Figure 3C, 4B) but no other branches.

In striking contrast to the limited number of *Gcna* and *mh* duplication events across *Drosophila*, we uncovered a staggering 15 *CG2694* duplications (Figure 3D, 4). Consistent with previous results, we found that the *CG2694* retrogene, *CG11322*, is restricted to the *melanogaster* subgroup ((Dai, et al. 2006; Zhang, et al. 2010); Figure 4A). We identified five additional *CG2694* duplication events in the *melanogaster* group (Figure 4A). Beyond the *melanogaster* group, we detected eight *CG2694* duplications in the *obscura* group (Figure 4B) and one duplication event in the *virilis-repleta* group (Figure 4C). Virtually all of the species that we assayed have at least one *CG2694* duplicate, though the identity of that young genes turns over in a revolving door pattern of gene family evolution (Demuth and Hahn 2009; Levine, et al. 2012). Moreover, almost all young *CG2694*-derived genes are autosomal, while *CG2694* is *X*-linked. An intriguing exception to this *X*-to-autosome directional gene movement was a large cluster of *CG2694* daughter genes dispersed in repetitive intergenic sequences across the *D. miranda X* chromosome (Figure 3D, 4B). Notably, the *Y* chromosome is only sparsely represented in some of the sampled genome assemblies and is entirely absent from others; consequently, we were unable to systematically assay gene duplication on the *Y* chromosome.

**Figure 4.**
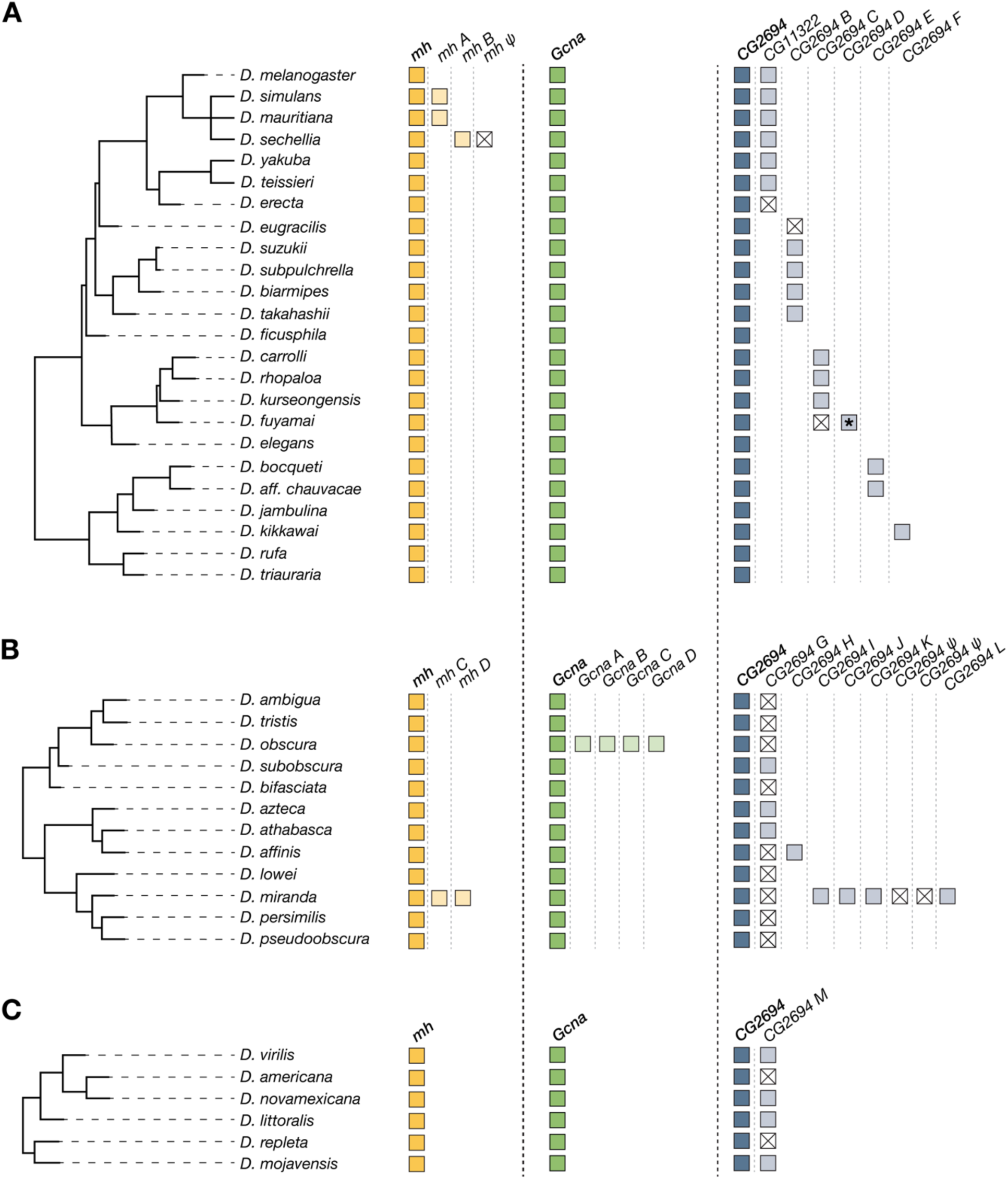
Pervasive gene birth and death across the Drosophila Spartan gene family. Summary of gene presence/absence across species trees of the (**A**) *melanogaster*, (**B**) *obscura*, and (**C**) *virilis-repleta* clades. Paralogs of *mh* (yellow), *Gcna* (green), and *CG2694* (blue) are indicated by boxes with lighter shades. *CG2694 C* in *D. fuyamai* (*) does not encode a SprT domain. Detectable pseudogenes are indicated by a white box with an “X”.

### Recurrent evolution of male germline-restricted expression of young Spartan family paralogs

In *D. melanogaster, mh and Gcna* are expressed primarily in the female germline (Delabaere, et al. 2014; Tang, et al. 2017; Bhargava, et al. 2020), *CG2694* is expressed in both the male and female germlines, and the young *CG2694* duplicate, *CG11322*, is expressed in the testis (Dai, et al. 2006; Zhang, et al. 2010). The divergent expression pattern between *CG2694* and its daughter gene, *CG11322*, raises the possibility that Spartan expression may diversify recurrently, mirroring the rapid evolution of the underlying DNA sequence. To test this idea, we focused on a subset of species that encode distinct suites of Spartan family genes. Consistent with previous reports, we found that Spartan family proteases are primarily expressed in the germline (Dai, et al. 2006; Zhang, et al. 2010; Delabaere, et al. 2014; Tang, et al. 2017; Bhargava, et al. 2020), although we also detect some low-level somatic expression (Figure 5). The ovary-biased expression pattern of the slowly-evolving *Gcna* is largely conserved, while *mh* undergoes transitions between ovary-biased, ovary- and testis-biased, and testis-restricted.

**Figure 5.**
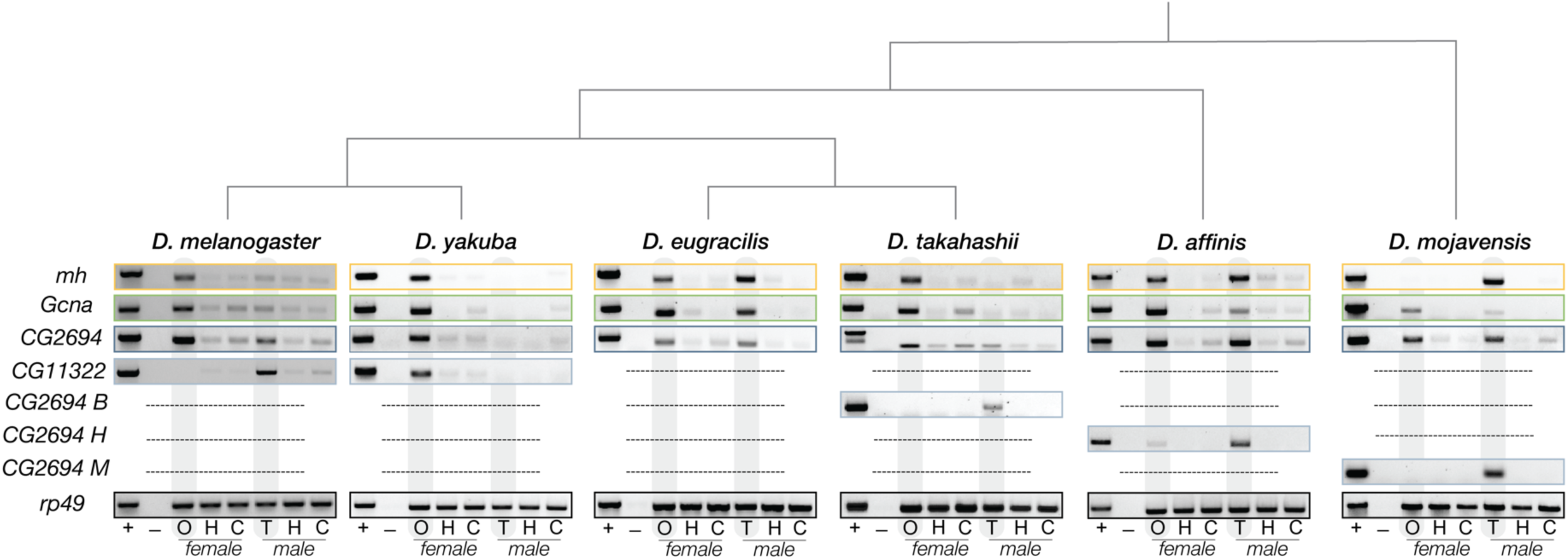
Parent-daughter gene divergence in tissue-specific expression profiles across Spartan family. RT-PCR products of Spartan family genes amplified across a panel of adult tissues (+ = genomic DNA positive control, - = water negative control, O = ovaries, H = head, C =carcass, and T = testis). PCR of template prepared without reverse transcriptase confirmed the absence of genomic DNA contamination (Figure S4).

The fast evolving and highly prolific *CG2694* shows a conserved ovary and testis expression profile across orthologs but dramatic differences in expression between parent and daughter genes. A previous report that a young *CG2694* duplicate is restricted to the testis (Dai, et al. 2006; Zhang, et al. 2010) was recapitulated in three, independently evolved *CG2694* young duplicates (Figure 5). This finding indicates that young, diverged *CG2694* duplicates repeatedly lose the ovary expression of their parent gene but evolve or retain testis expression. Combined with a previously reported testis-restricted *mh* duplicate in *D. simulans* (Chakraborty, et al. 2021), these data suggest that DNA-protein crosslink repair may require recurrent innovation, or at least specialization, in the male germline. A striking exception to this expression pattern is found in *D. yakuba*, where not one Spartan family gene is expressed in the testis (Figure 5).

This evidence of Drosophila-wide expression evolution across the Spartan family, combined with recurrent duplication and adaptive sequence evolution across multiple timescales and across multiple genes, implicates an ever-present agent of selection driving innovation of DNA-protein crosslink repair.

### A model of antagonistic coevolution between Spartan proteins and satellite DNA

What is the agent of selection driving Spartan family diversification? Published evidence of antagonistic coevolution between *mh* and a DNA satellite array implicates DNA repeats (Brand and Levine 2022). The *D. simulans* version of *mh* appears to over-actively cleave crosslinks between a *D. melanogaster*-specific satellite array and the essential enzyme, Topoisomerase 2. Intriguingly, the *D. simulans* genome has much lower repeat content compared to the *D. melanogaster* genome (5% vs 20% of the genome, respectively) (Lohe and Brutlag 1987), suggesting that repeat-poor genomes evolve highly active Spartan proteases and repeat-rich genomes evolve comparatively less active Spartan proteases.

Why might DNA satellite expansions and contractions trigger adaptive evolution of Spartan protease activity? Here we propose a model under which the secondary structures enriched at DNA satellites are the primary drivers of Spartan family evolution. Recent biochemical analysis of human Spartan revealed that single-/double-stranded DNA junctions (“ss/dsDNA junctions”) promote Spartan protease activity (Reinking, et al. 2020). These ss/dsDNA junctions are present not only at replication forks that recruit Spartan (Juhasz, et al. 2012) but also at DNA secondary structures and RNA/DNA hybrids (Zheng and Dean 2009; Loomis, et al. 2014; Velazquez Camacho, et al. 2017; Xu, et al. 2017). Importantly, these DNA secondary structures and RNA/DNA hybrids, as well as stalled replication forks, are enriched at satellite DNA (Nadel, et al. 2015; Bacolla, et al. 2016; Madireddy and Gerhardt 2017; Polleys, et al. 2017; Velazquez Camacho, et al. 2017); Figure 6A). Satellite expansion would increase ss/dsDNA junction abundance while satellite contraction would reduce ss/dsDNA junctions. Elevated ss/dsDNA junctions may tip the balance of Spartan activity toward too much proteolysis while reduction of ss/dsDNA junctions may tip the balance towards too little. To restore healthy rates of DNA-protein crosslink repair, regulation of the local Spartan protein dose may evolve though multiple mechanisms. Changes to transcript and/or protein abundance, protein affinity for the replication fork, and/or self-cleavage of excess Spartan protein (Perry and Ghosal 2022) can each alter Spartan protein dose and rebalance DNA-protein crosslink repair rates.

**Figure 6.**
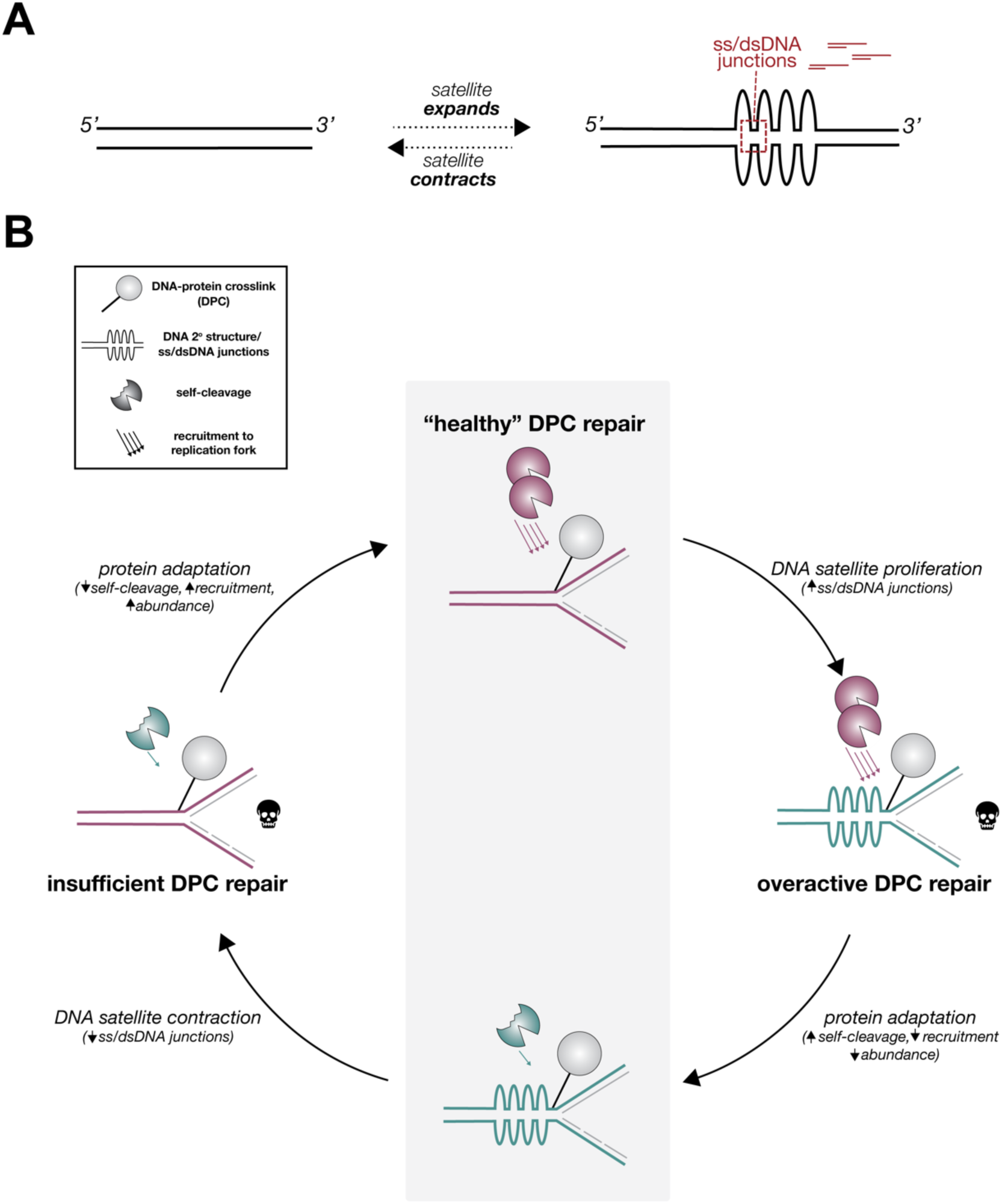
A seesaw model of DNA satellite–Spartan family coevolution accounts for pervasive Spartan family evolution. (**A**) Fluctuations in DNA satellite copy number change the abundance of ss/dsDNA junctions linked to secondary structures and RNA/DNA hybrids. DNA satellite proliferation results in an increased abundance of ss/dsDNA junctions while DNA satellite contraction results in decreased ss/dsDNA junctions. (**B**) The seesaw model is predicated on the idea that Spartan protease activity is altered by 1) changes in ss/dsDNA abundance linked to DNA satellite fluctuations, 2) Spartan affinity for the replication fork, 3) the rate of Spartan self-cleavage, and 4) the abundance of Spartan protein. When intrinsic Spartan protease regulation balances the abundance of ss/dsDNA junctions, a healthy rate of DNA-protein crosslink (DPC) repair is sustained (center). Over evolutionary time, DNA satellite proliferation and contraction perturbs overall protease activity. “Mismatched” protein and satellite DNA (aqua and magenta) result in overactive DPC repair (right) or insufficient DPC repair (left). These deleterious consequences of satellite fluctuations trigger protein adaptation that either increases or decreases protein abundance, recruitment to chromatin, and/or Spartan protease self-cleavage to restore healthy DPC repair rates.

We propose that the preservation of healthy DNA protein-crosslink repair in the wake of DNA satellite expansion/contraction drives a coevolutionary “seesaw.” Under this model, adaptive evolution of intrinsic Spartan protease regulation – protein abundance, affinity to the replication fork, and/or strength of self-cleavage – balances the fluctuating abundance of ss/dsDNA junctions that also modulate proteolysis. This balance maintains healthy levels of DNA-protein crosslink (“DPC”) repair. Over evolutionary time (Figure 6B), elevated ss/dsDNA junctions linked to DNA satellite proliferation trigger overactive DNA-protein crosslink repair that prematurely cleaves crosslinks required for essential DNA-templated functions. The fitness reduction associated with excess Spartan protease activity selects for reduced protein abundance, reduced replication fork recruitment, and/or increased self-cleavage to restore healthy rates of DNA-protein crosslink repair. DNA satellite contraction then reduces ss/dsDNA abundance, leading to insufficient Spartan activity and persistent crosslinks that imperil replication and transcription. The fitness reduction associated with deficient Spartan protease activity triggers selection for increased protein abundance, increased replication fork recruitment, and/or decreased self-cleavage to restore healthy repair rates. This coevolutionary seesaw, driven by species-specific satellite expansions and contractions, would result in phylogeny-wide sequence and expression diversification of Spartan family genes.

The seesaw model of DNA satellite-Spartan protease coevolution specifically predicts that satellite DNA turnover shapes Spartan regulation. Spartan regulation maps to C-terminal domains that mediate affinity to the replication fork and self-cleavage (Centore, et al. 2012; Lopez-Mosqueda, et al. 2016; Stingele, et al. 2016; Vaz, et al. 2016). Pervasive adaptive sequence evolution along both short evolutionary timescales (∼2.5my) and long evolutionary timescales (∼12-25my), across the ancient family members and between parent and young daughter genes, have the potential to alter Spartan activity. Moreover, domain-specific analysis of positive selection at *mh* in *D. melanogaster* and *D. simulans* revealed that adaptive substitutions are enriched in the replication fork recruitment domain and the regulatory domain of self-cleavage (Brand and Levine 2022). Three additional species pairs, sampled from each of the three focal Drosophila clades, recapitulate this pattern of elevated *dN/dS* across the C-terminus (Figure S5). Modulating local protein dose at the replication fork (or another genome structure) can also be achieved by expression evolution. Indeed, we uncovered many examples of minimal or no expression in either the soma or germline tissue of one species but high expression in these same tissues in a closely related species. Biochemical analysis of Spartan proteases derived from species with exceptionally high or low amounts of DNA repeats would be a natural next step to testing this seesaw model of DNA satellite-Spartan family coevolution.

The seesaw model accounts for Spartan family gene expression evolution over time; however, the model does not account for the recurrent evolution of male-germline restricted young duplicate genes. The enrichment of young genes expressed in the testis implicates several possible evolutionary pressures: escape from *X*-linked gene silencing in the male germline, sexual antagonism, and intra-genomic conflict. *D. melanogaster* hyperactivates the *X* chromosome to dosage compensate in the heterogametic male. The expression of *X*-linked genes is considerably lower than autosomal genes in the testis (Landeen et al. 2016). This failure to dosage compensate in the testis can promote *X* to autosome duplication events to escape reduced expression (Betran, et al. 2002; Zhang, et al. 2010). Consistent with this framework, *CG2694* is *X*-linked while nearly all of its young duplicates are located on autosomes (Table S3) and many have testes-restricted expression (Figure 5). Sexual antagonism might also drive this testis bias. The male and female germline may have different, and opposing, requirements for DNA-protein crosslink repair. For example, the ovary-specific Spo11 enzyme crosslinks to DNA as it creates double-strand breaks that promote meiotic recombination in the Drosophila female germline (Keeney, et al. 1997; McKim and Hayashi-Hagihara 1998). A young testis-specific protease can act independently of such ovary-specific functions. In addition to escape from *X*-silencing and sexual antagonism, intra-genomic conflict may drive the testes expression bias of young duplicates. A selfish, *X*-linked Spartan family gene could bias its transmission to the next generation by avoiding repair of crosslinks on the DNA satellite-rich *Y* chromosome, rendering *Y*-bearing sperm (or embryos fertilized by that sperm) inviable. These males would produce female-biased progeny, triggering the evolution of autosomal and *Y*-linked suppressors, that typically arise via duplications of the driver, to restore equal sex ratio (Courret, et al. 2019). Regardless of the evolutionary pressure, our data suggest that the Spartan protease activity in the male germline is acutely sensitive to DNA satellite fluctuations.

The notion that DNA satellite contractions and expansions can shape fundamental cellular processes challenges the historical assumption that repeat-rich genomic regions are largely junk (Ohno 1972). DNA-protein crosslink repair now joins an ever-growing list of vital biological functions impacted by DNA satellites, including chromosome segregation, telomere integrity, sperm maturation, and nuclear architecture (Schoeftner and Blasco 2009; McKinley and Cheeseman 2016; Iwata-Otsubo, et al. 2017; Jagannathan, et al. 2018; Mills, et al. 2019).

However, our grasp of how and why repeat evolution might perturb important cellular processes and development is still in its infancy (Brand and Levine 2021). Excitingly, the list of complete, telomere-to-telomere genome assemblies is growing rapidly (Liu, et al. 2020; Gonzalez de la Rosa, et al. 2021; Rhie, et al. 2021; Nurk, et al. 2022; Huang, et al. 2023; Kurokochi, et al. 2023; O’Donnell, et al. 2023), offering unprecedented opportunities to study the impact of DNA satellite evolution on health, disease, and the formation of new species.

## MATERIALS AND METHODS

### Population genetic and molecular evolution analyses

We conducted population genetic analysis of Spartan family genes using polymorphism and divergence data from *D. melanogaster* and *D. simulans*. We extracted nine *D. melanogaster* alleles of *Gcna* (*X*:1862009-1865835), *CG2694* (*X*:2714079-2717657), and *CG11322* (*2L*:6757864-6759628) from genome assemblies of lines collected from Lyon, France (dmel r6.4). For each gene, we also extracted 10 *D. simulans* alleles from genome sequencing reads generated from lines collected from Nairobi, Kenya. We used bowtie2 (Langmead and Salzberg 2012) to map short-read Illumina reads to a long-read based *D. simulans* reference genome (Chang, et al. 2022). We called variants using bcftools v1.8 (Li 2011), considering only reads with mapping quality >20 (*i.e.*, unique-mapping reads) and only polymorphic sites for which there were at least ten reads supporting the SNP. We aligned the sequences in Geneious using the Geneious Alignment algorithm under default settings (Geneious v11.1.5, Biomatters, Auckland, New Zealand) and confirmed alignment quality by eye. We performed a McDonald-Kreitman test (McDonald and Kreitman 1991) with the *D. melanogaster* and *D. simulans Gcna*, *CG2694*, and *CG11322* coding sequences. To assess lineage-specific evolution of *CG2694* and *CG1132,* we performed a polarized McDonald-Kreitman test using *D. yakuba* as an outgroup. We previously reported the results of both pairwise and polarized McDonald-Kreitman tests for *mh* (Brand and Levine 2022).

To test for evidence of recurrent adaptive evolution over longer evolutionary timescales, we conducted a phylogeny-based molecular evolution analysis of Spartan family orthologs. To avoid synonymous site saturation, we conducted these analyses in three distinct clades that each span ∼10-25 million years of evolution: the *melanogaster* group, the *obscura* group, and the *virilis-repleta* group (Table S2). Once we compiled the coding sequences of each gene across all 42 focal species (see below for ortholog identification methods), we aligned the amino acid sequences using the Geneious Translation Alignment algorithm. We visually assessed and gap-adjusted the coding sequence alignments to retain in-frame codons. For each gene, we fit our multiple alignments to an NSsites model, which is part of the CODEML package in PAML (Yang 1997). To determine statistical significance, we used a likelihood ratio test that compared model 7 (*dN/dS* values fit a beta distribution from 0 to 1) and model 8 (model 7 parameters and *dN/dS* > 1).

We calculated pairwise *dN/dS* between *mh* alleles using a window size of 200bp (step size = 20bp) in the software package, DnaSP (Librado and Rozas 2009).

### Identification of Spartan family orthologs and paralogs

To identify Spartan family gene paralogs, we conducted a tBLASTn search of 42 highly contiguous genome assemblies that span the *melanogaster*, *obscura*, and *virilis-repleta* clades (Table S2). We used the protein sequences of each ancient Spartan family gene found in *D. melanogaster*, *D. obscura*, *D. pseudoobcura*, and *D. virilis* (*mh*, *GCNA*, *CG2694*) as a query.

For each species, we extracted all hits with an e-value < 0.01. We aligned the SprT domain nucleotide sequence (Table S3) using MAFFT v7.401 with default parameters (Katoh and Standley 2013). We then cleaned the alignment by removing small gaps and ambiguously aligned codons using Gblocks v0.91.1 (Talavera and Castresana 2007). We built phylogenetic trees using maximum likelihood methods implemented in PhyML with a Smart Model Selection v1.8.1 (Guindon, et al. 2010; Lefort, et al. 2017). We calculated branch support using an approximate likelihood ratio test (Anisimova, et al. 2011). We also subjected all hits to synteny analysis to determine paralogs from orthologs, and to confirm orthologs across different genomes. We called a hit a pseudogene if we detected no in-frame start codon within 50 base pairs up- or downstream of the original start codon, or if a premature stop codon truncated >95% of the protein (three instances).

### Tissue-specific gene expression analysis

To determine the expression pattern of Spartan family orthologs and paralogs across adult tissues, we made cDNA from RNA prepared from the heads, ovaries, testes, and remaining carcasses of adult flies (Table S2). Specifically, we extracted RNA using the mirVana miRNA Isolation kit (Invitrogen, Waltham, MA) and then DNase-treated each sample (TURBO DNase, Invitrogen, Waltham, MA). We then prepared cDNA using the Superscript III First-strand Synthesis System (Invitrogen, Waltham, MA) including a “Reverse Transcriptase-minus” (RT-) control of each sample to rule out genomic contamination (Figure S4). To determine the tissue-specific expression pattern, we performed RT-PCR using gene- and species-specific PCR primers (Table S4) that amplify the Spartan family genes of interest. We used the ubiquitously expressed *rp49* gene as a positive control.

## ACKNOWLEDGEMENTS

We thank E. Marti for technical assistance with extracting *D. simulans* alleles and R. Unckless for kindly sharing fly stocks. We also thank the Levine Lab, L. Kursel, D. Dudka, and A. Das for feedback on the manuscript and the Levine Lab, M. Buszczak, and members of the Lampson Lab for vital discussions about the project. This work was supported by a Shurl and Kay Curci Foundation fellowship from the Life Sciences Research Foundation and NIH K99GM149943 to C.L.B. and NIH grant R35GM124684 to M.T.L.

## SUPPLEMENTARY FIGURE LEGENDS

**Figure S1.**
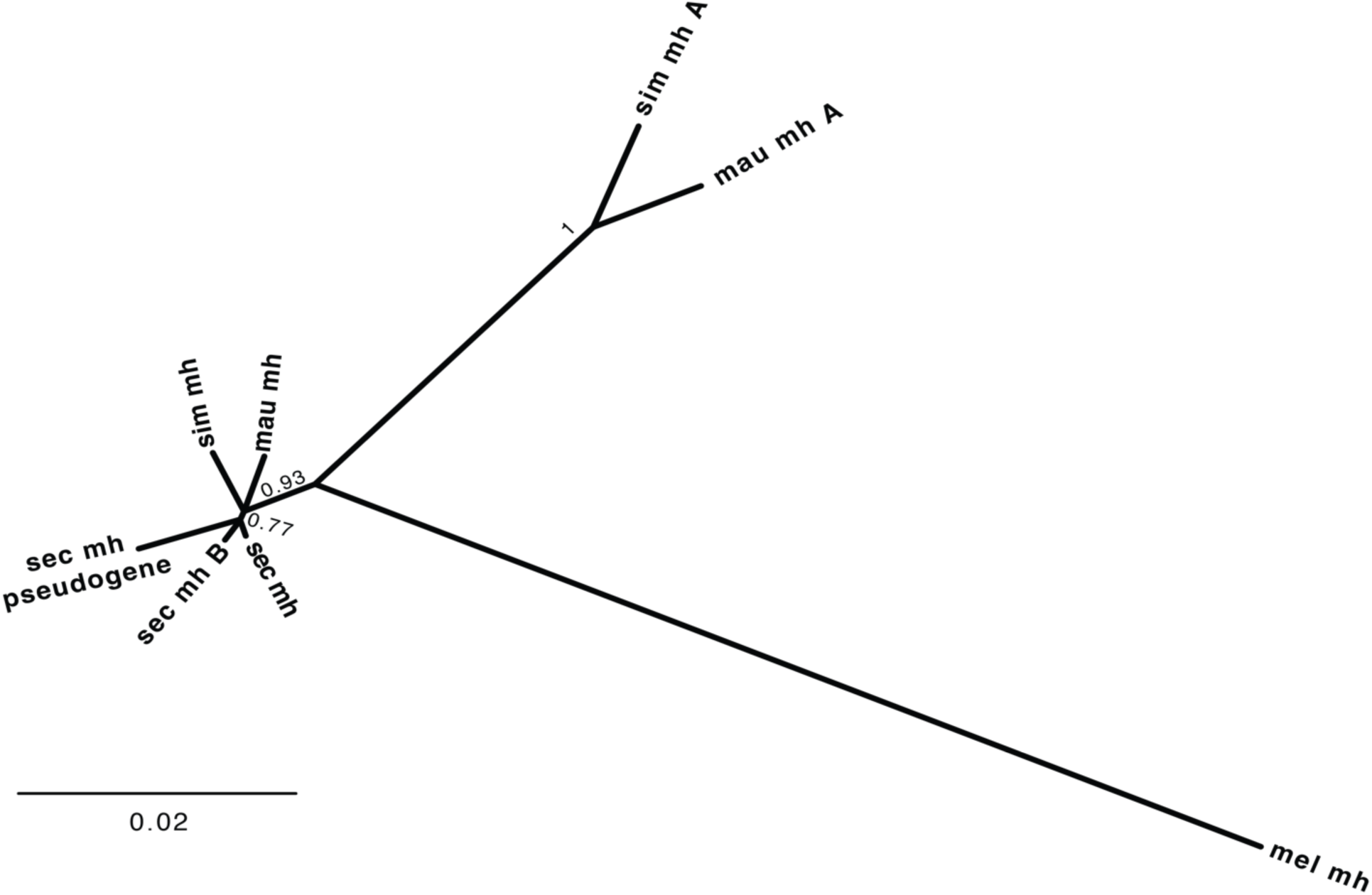
*mh* duplicated along the lineage leading to *D. sechellia*. The *D. sechellia mh-B* clusters away from the previously described *mh A* found in *D. simulans* and *D. mauritiana* (Chakraborty, et al. 2021). Only resolved nodes with branch support > 0.75 are shown.

**Figure S2.**
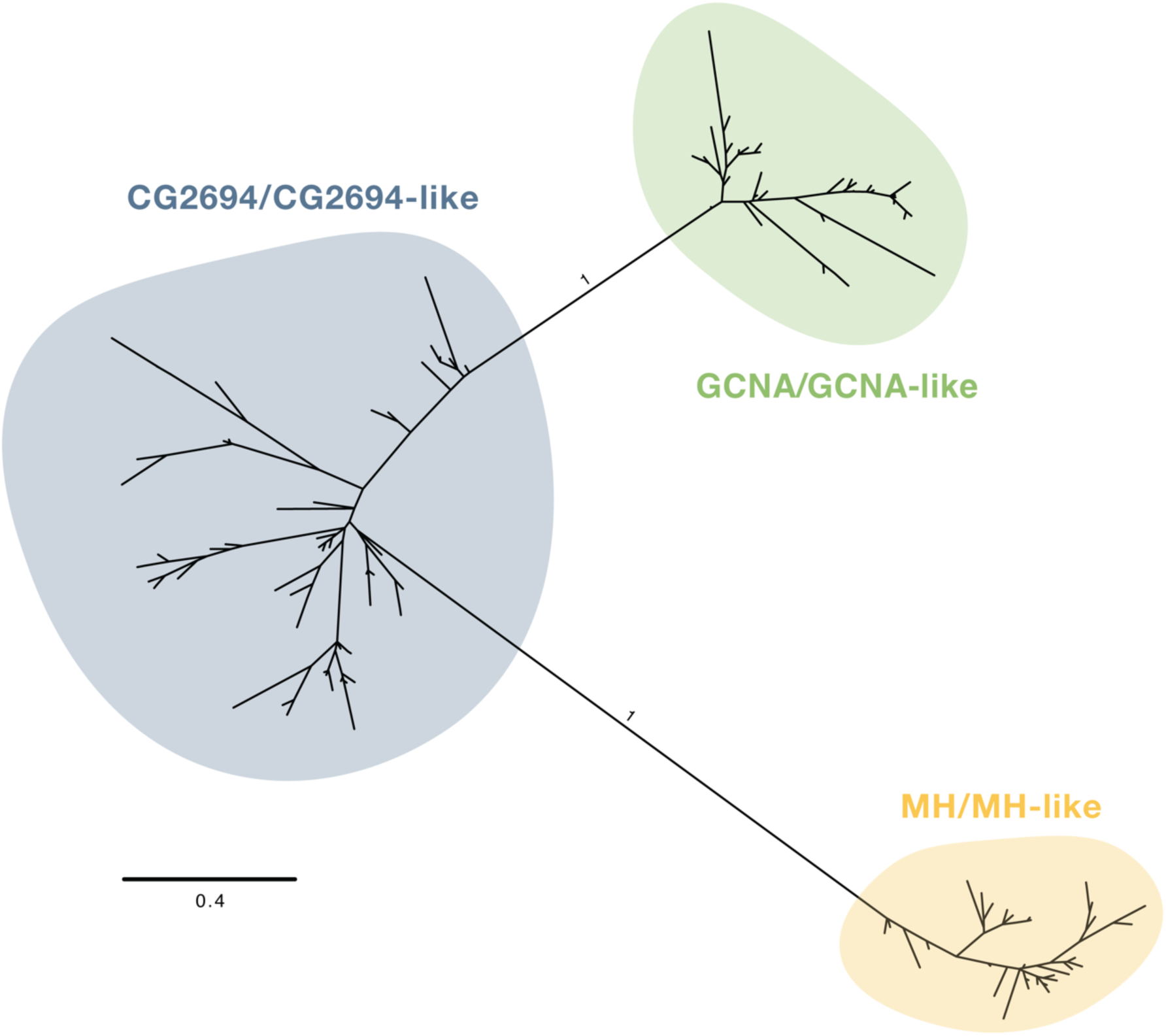
Phylogenetic tree of all significant hits and query sequences reveals three distinct Spartan subfamilies. Phylogenetic tree based on the SprT domain of the original query and all significant hits (Table S3) from the 42 focal genomes (Table S2). *Gcna, mh*, and *CG2694* subtrees can be found in Figure 3.

**Figure S3.**
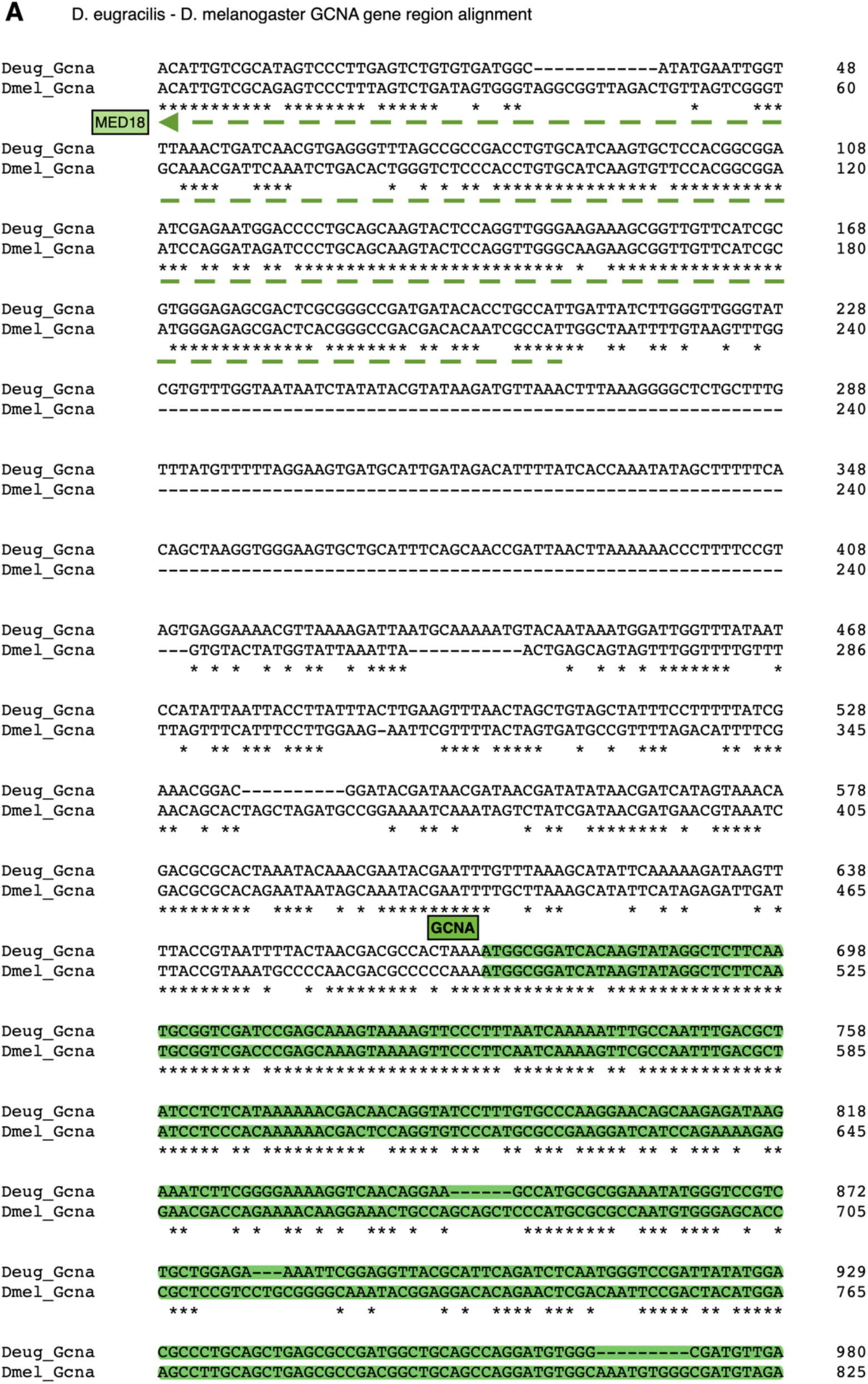
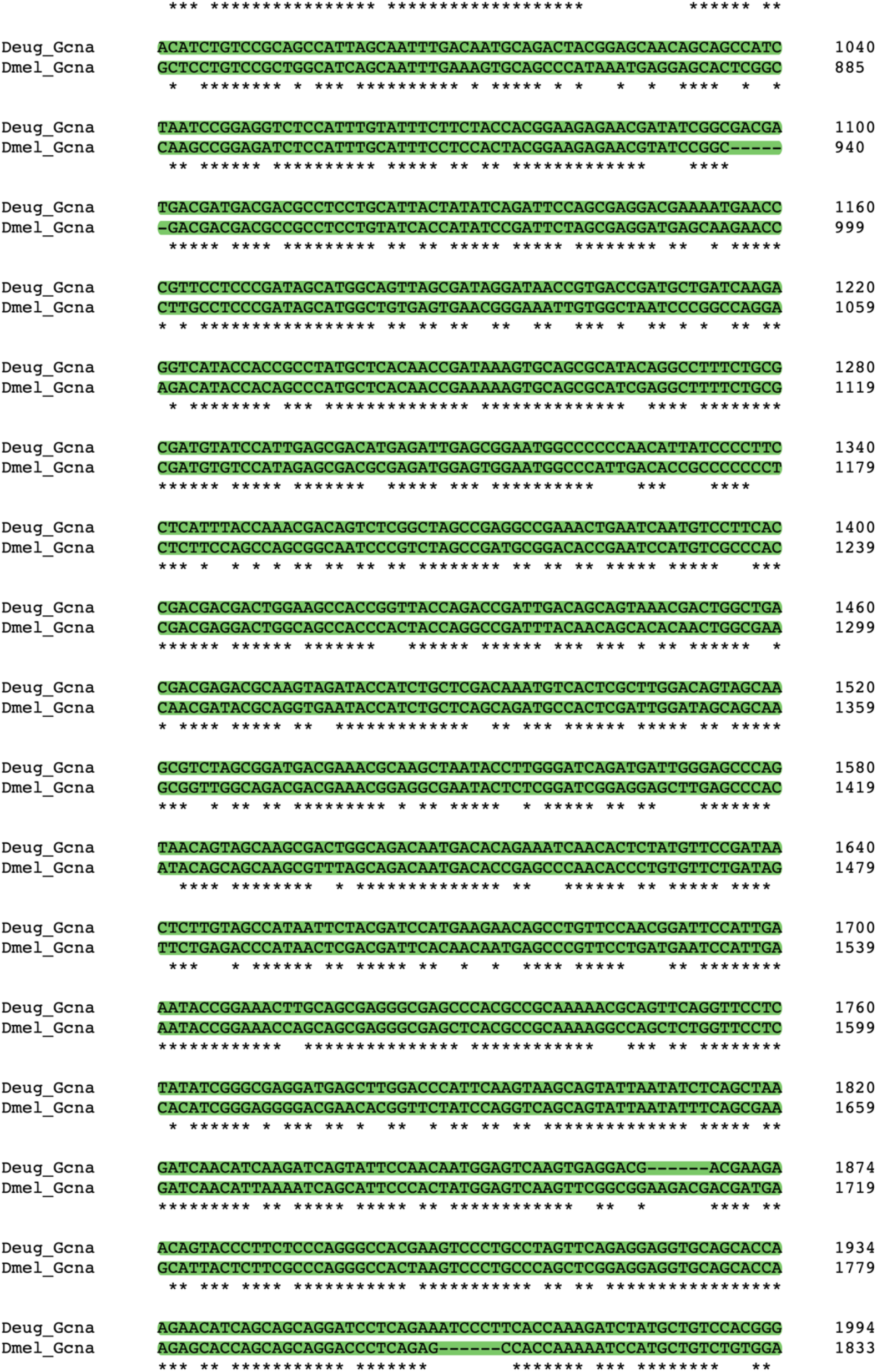
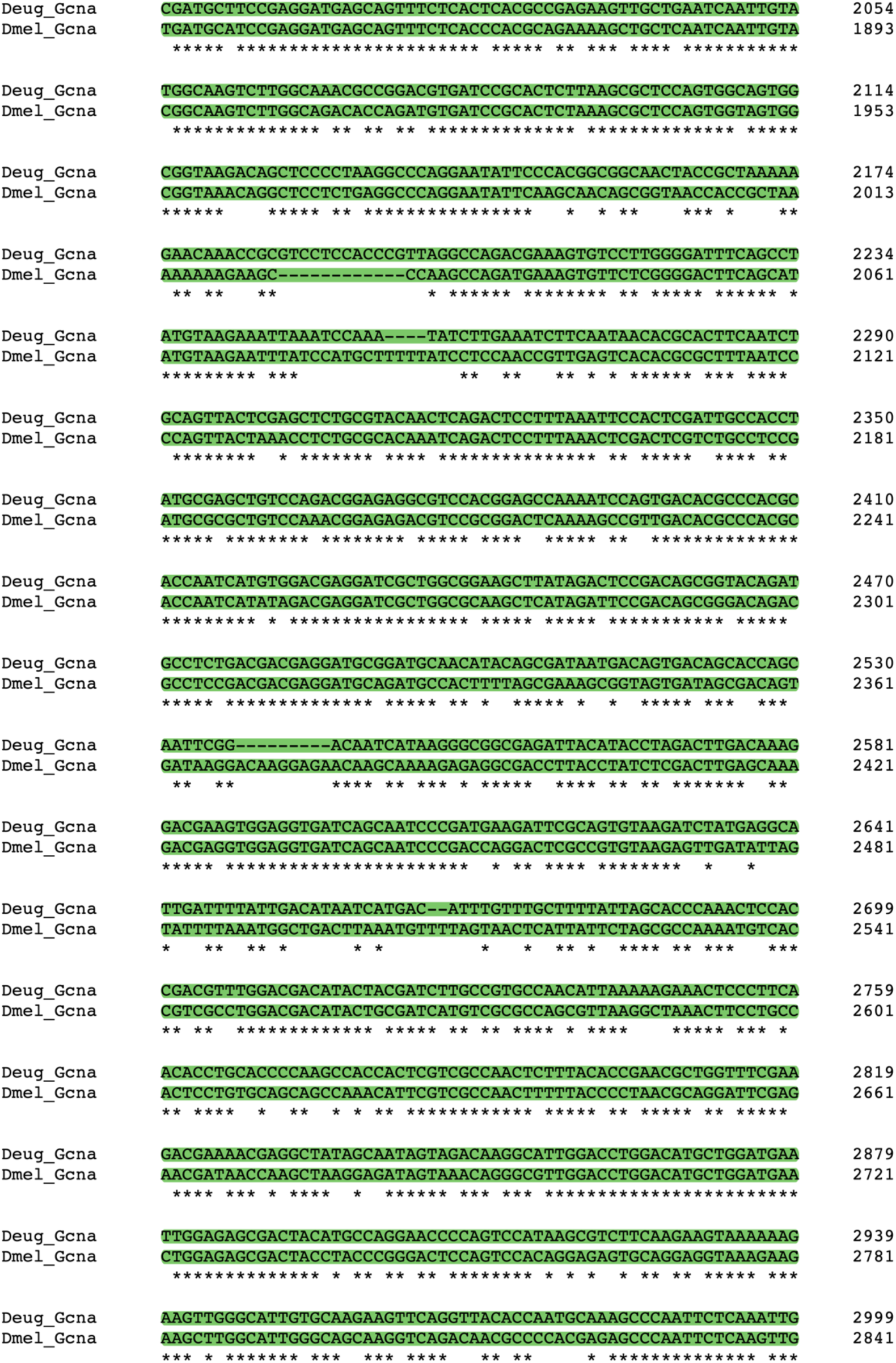
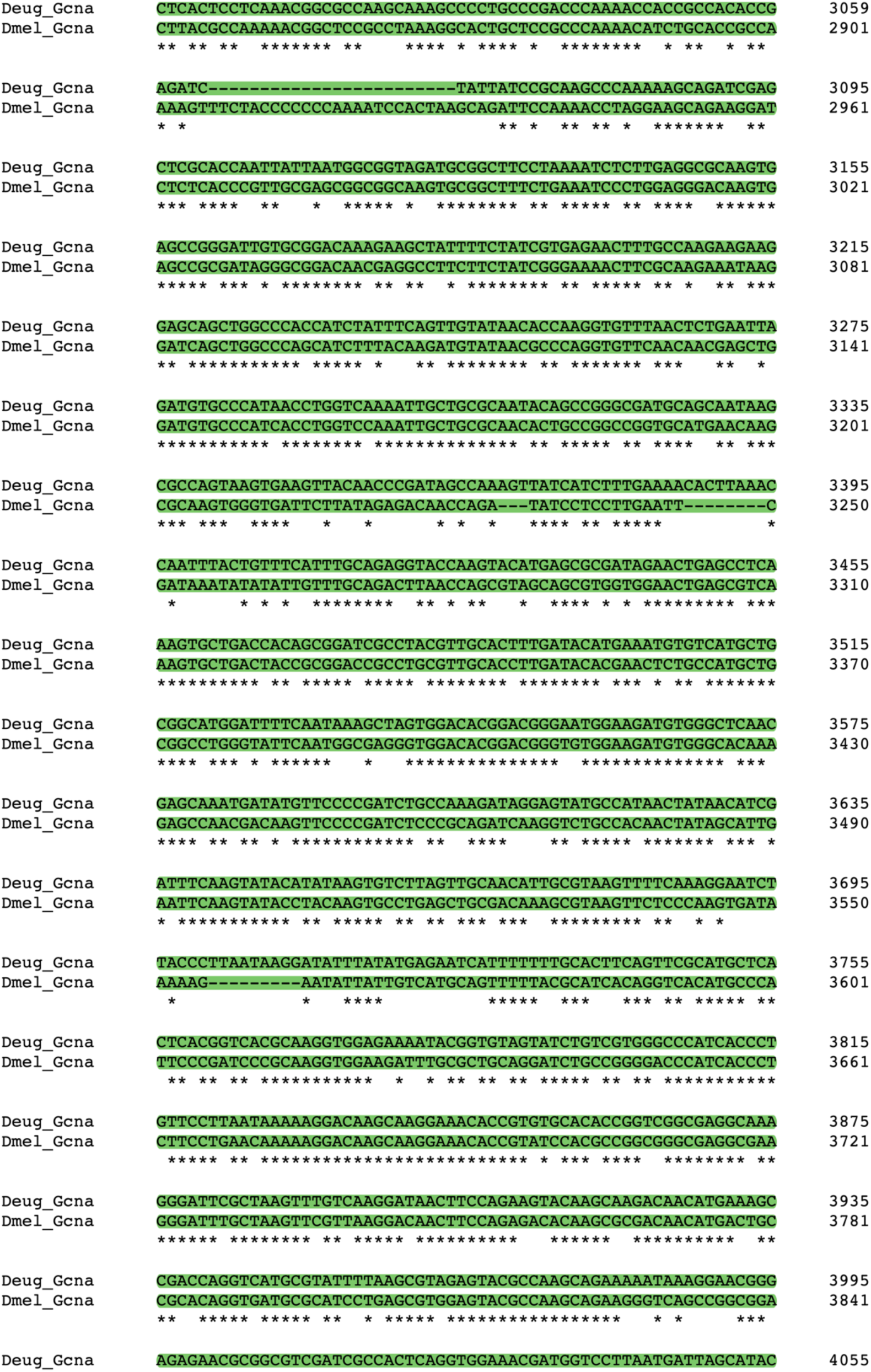
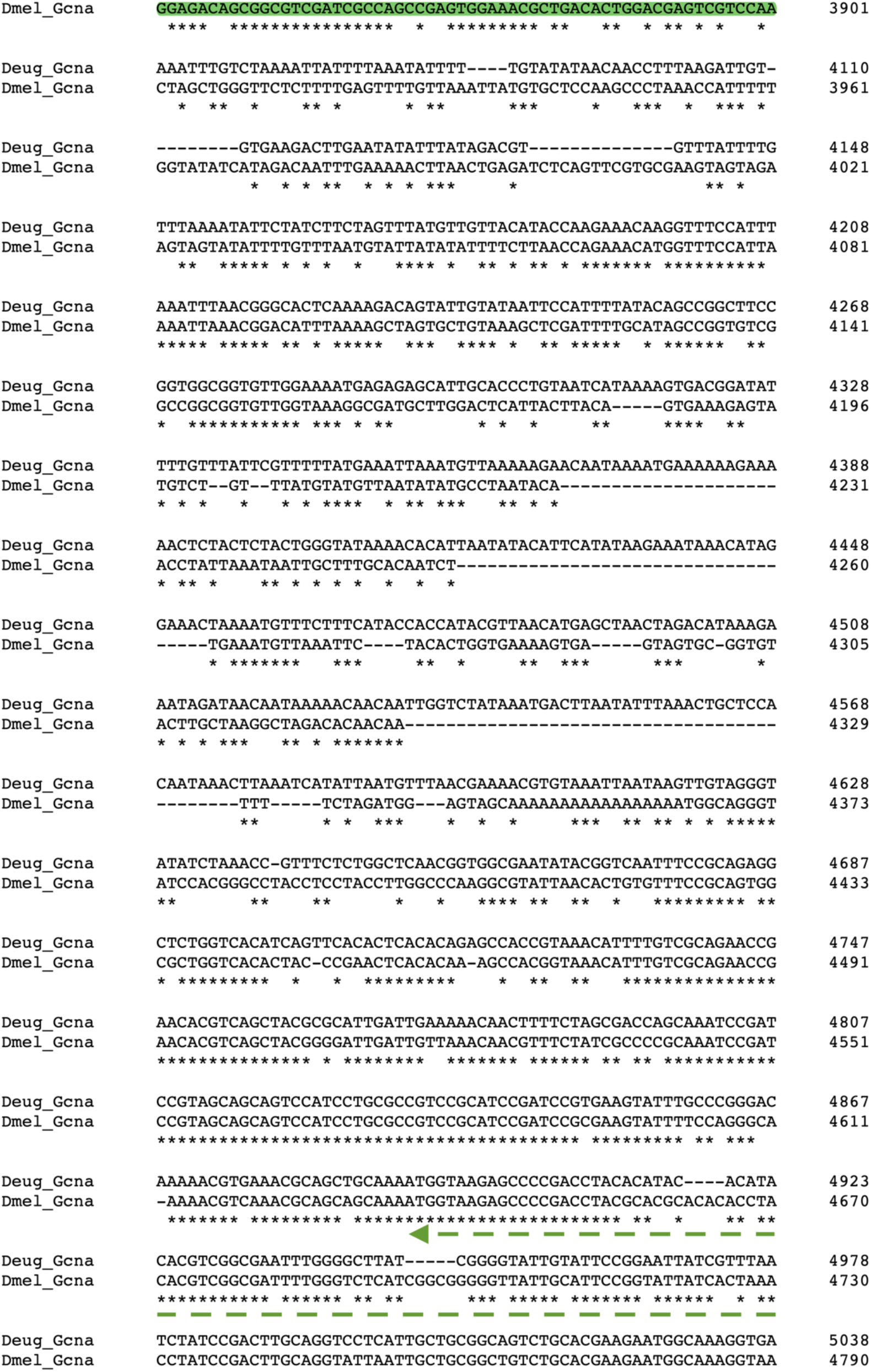
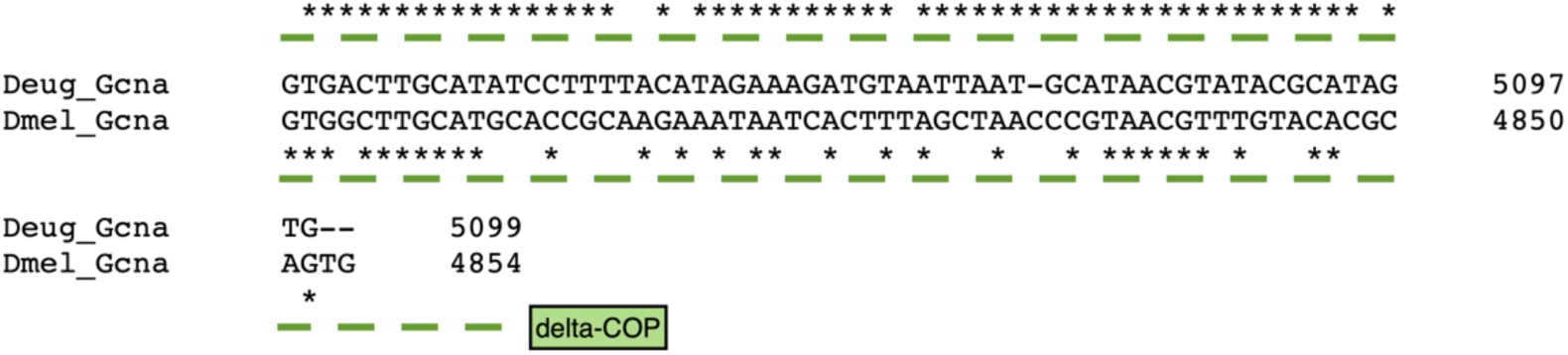
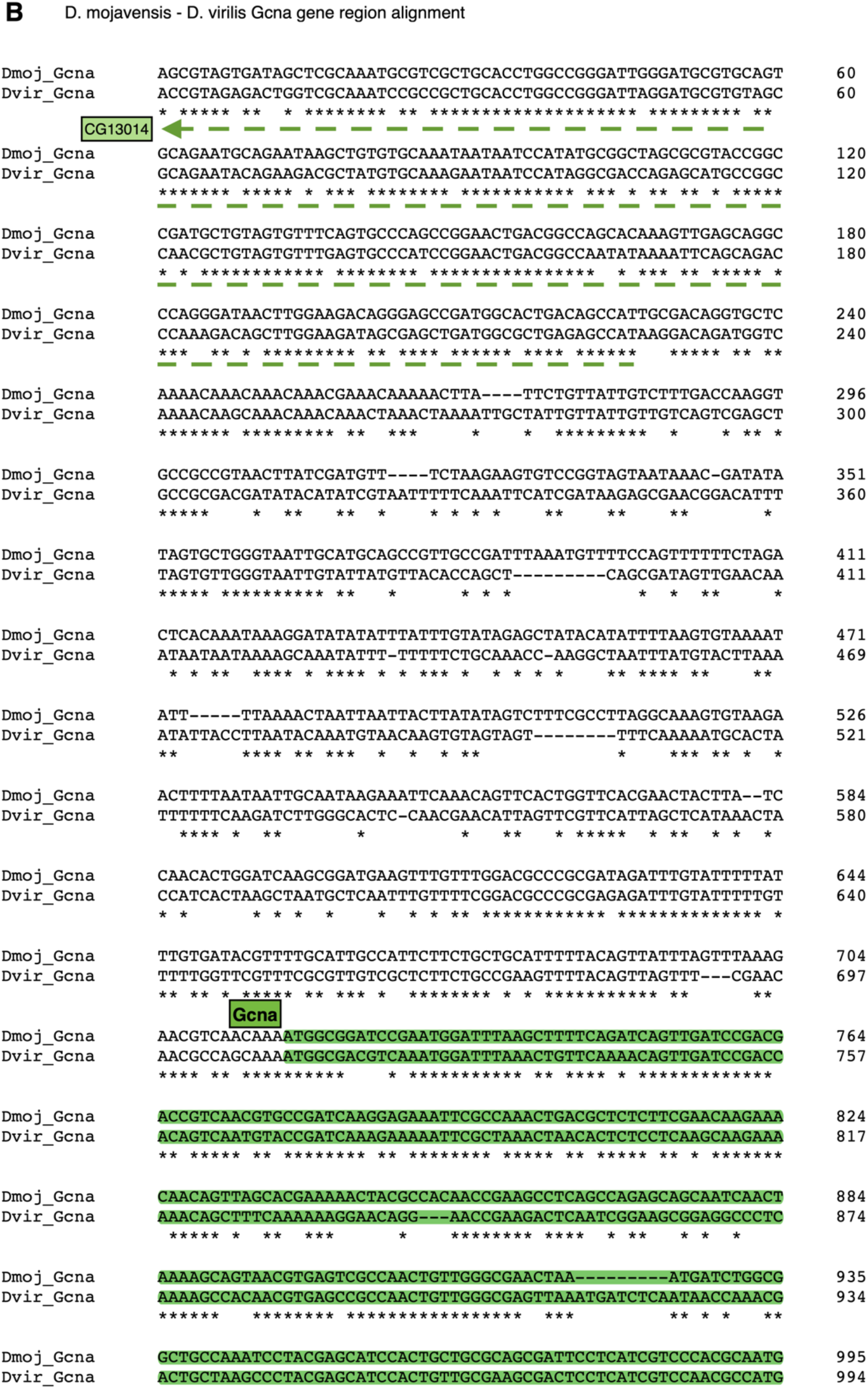
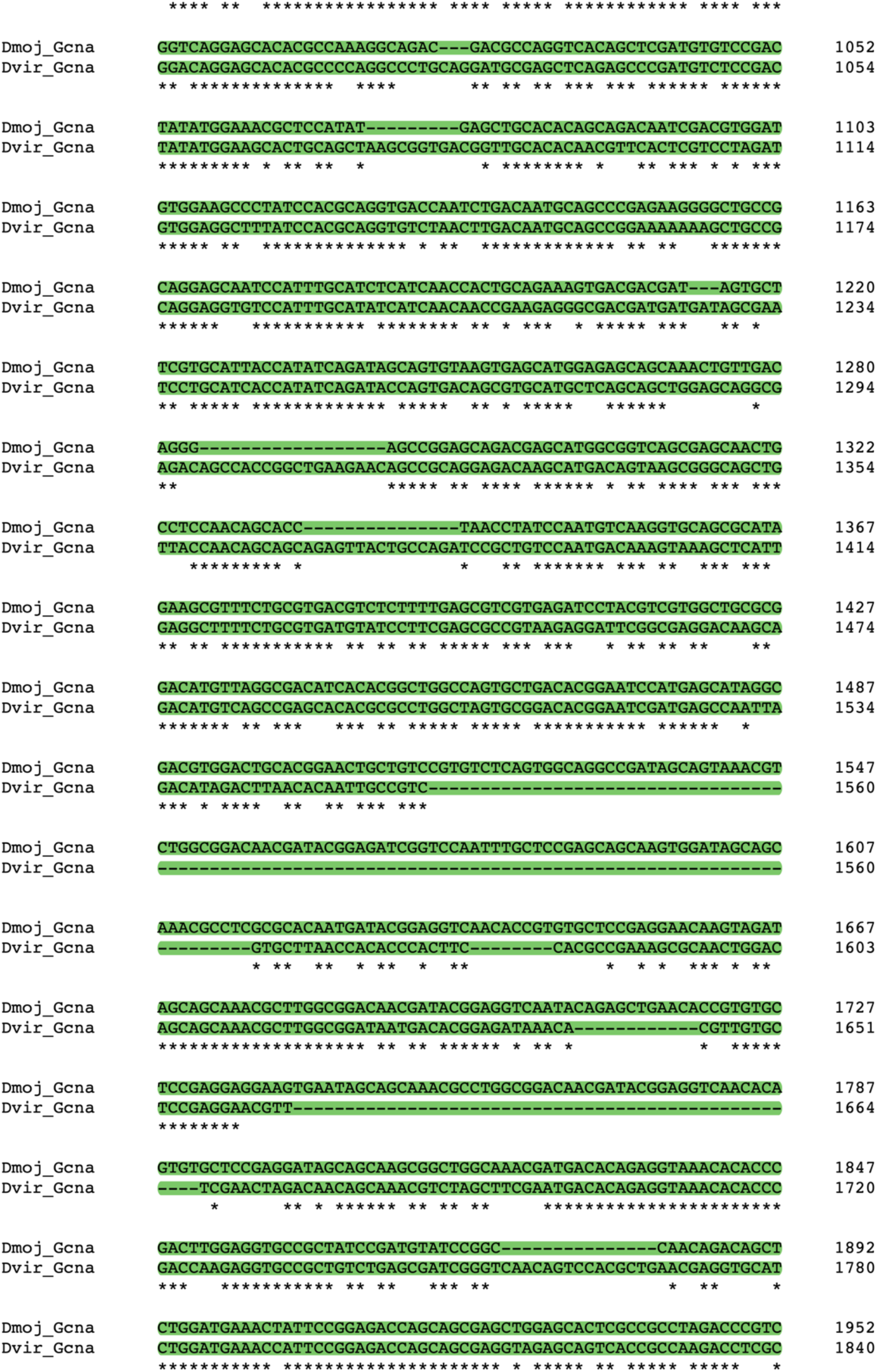
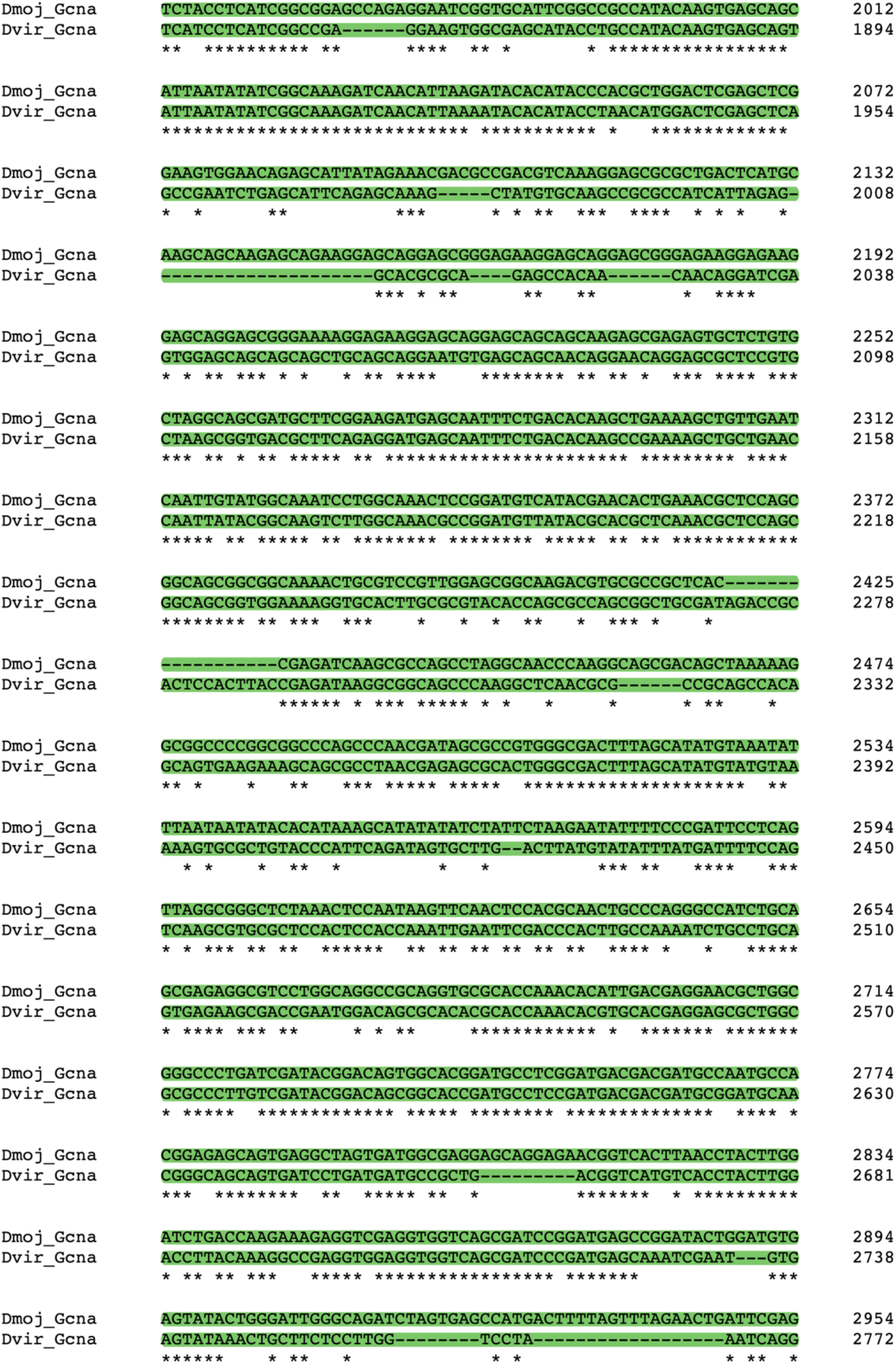
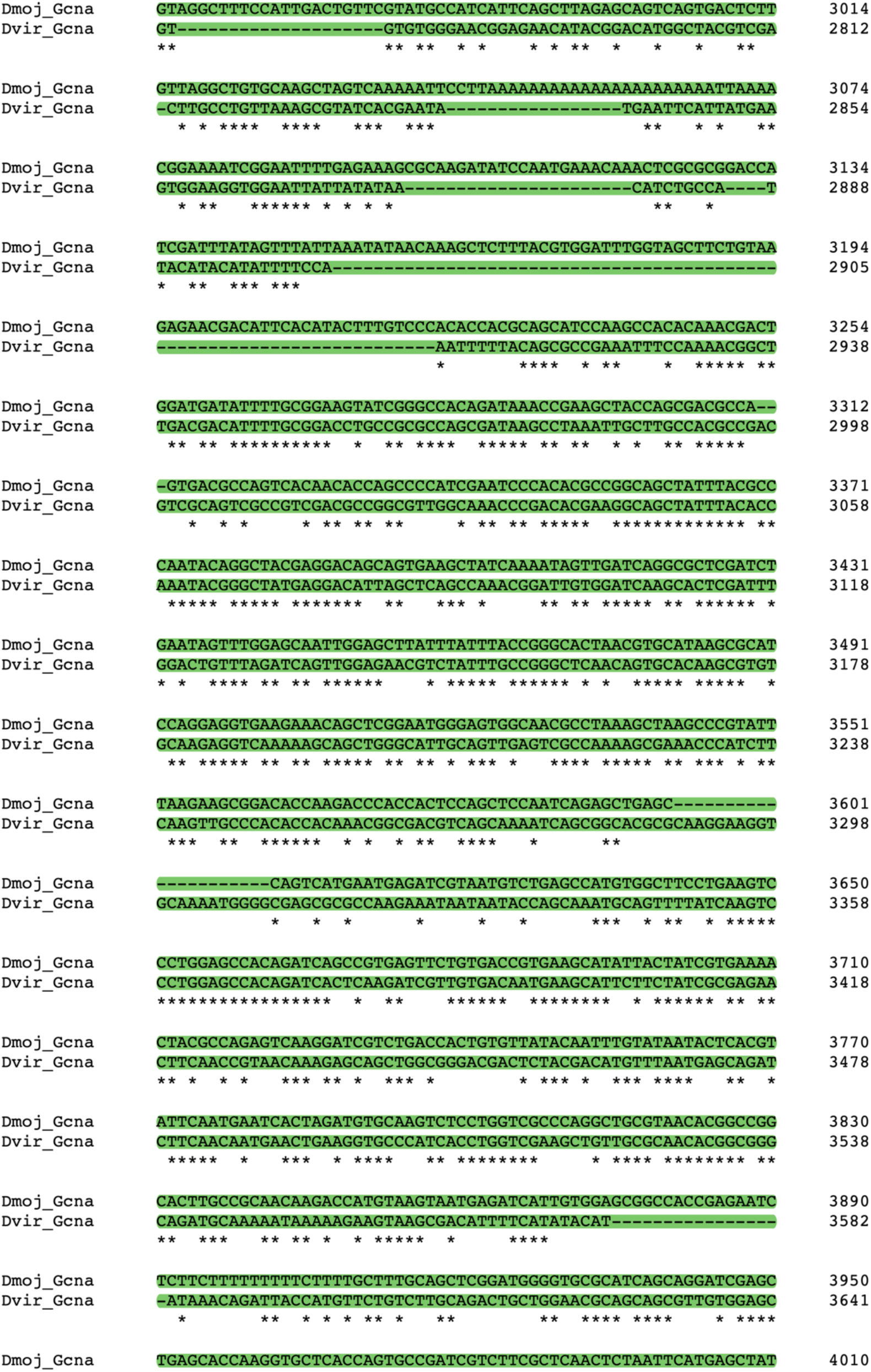
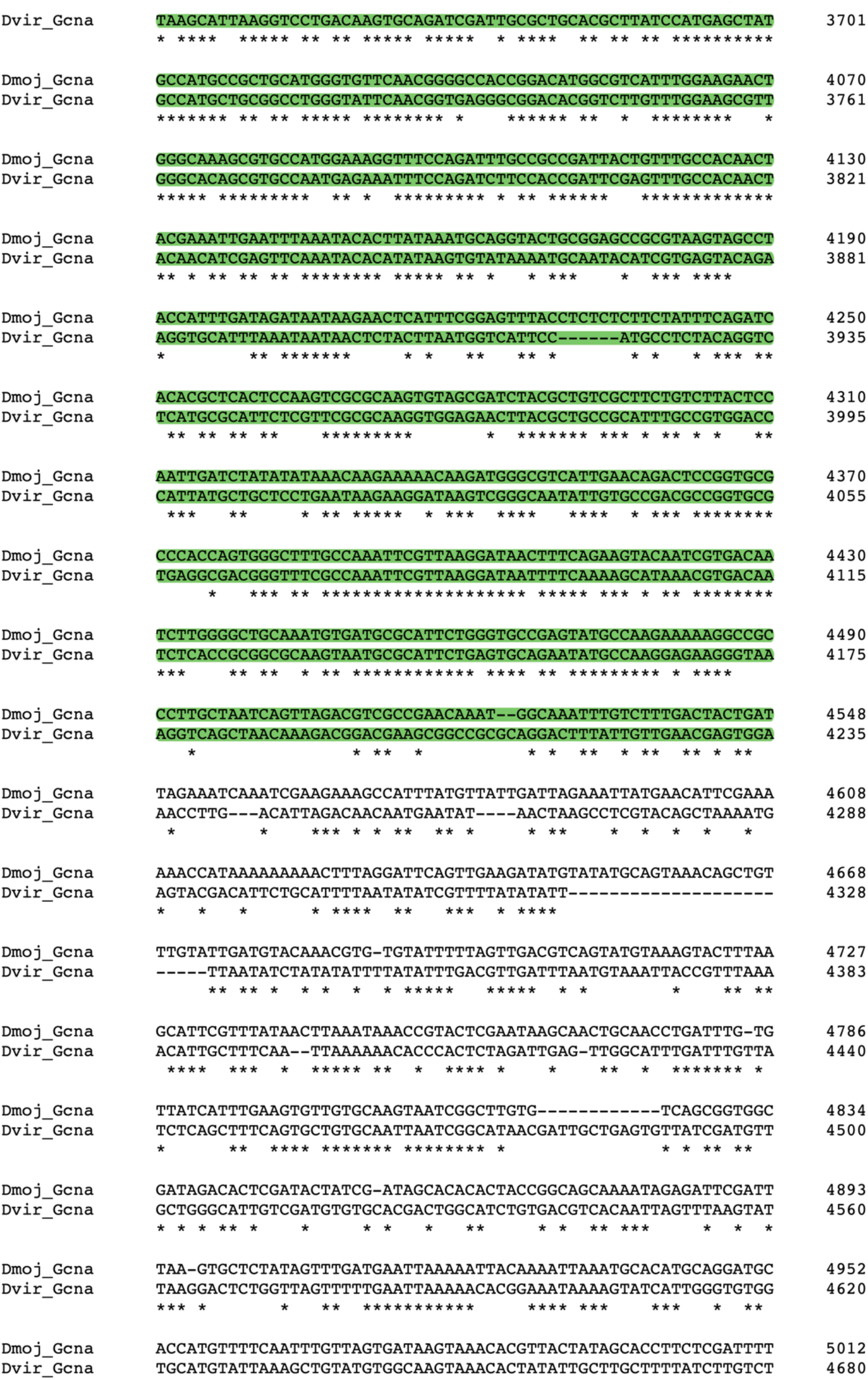
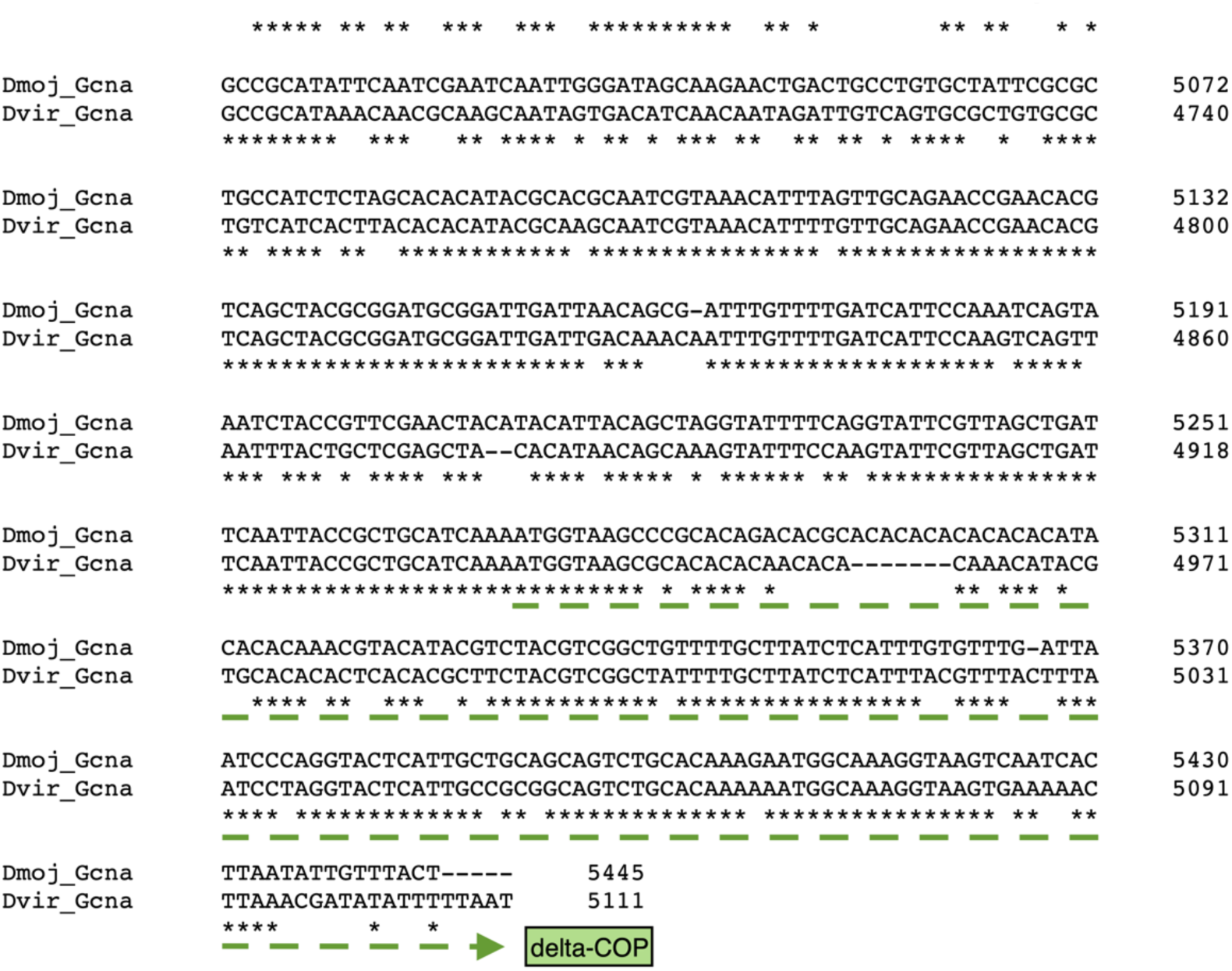
*D. eugracilis Gcna* and *D. mojavensis Gcna* are syntenic with *D. melanogaster Gcna*. Genes upstream and downstream of *Gcna* in each species are shown on the alignments.

**Figure S4.**
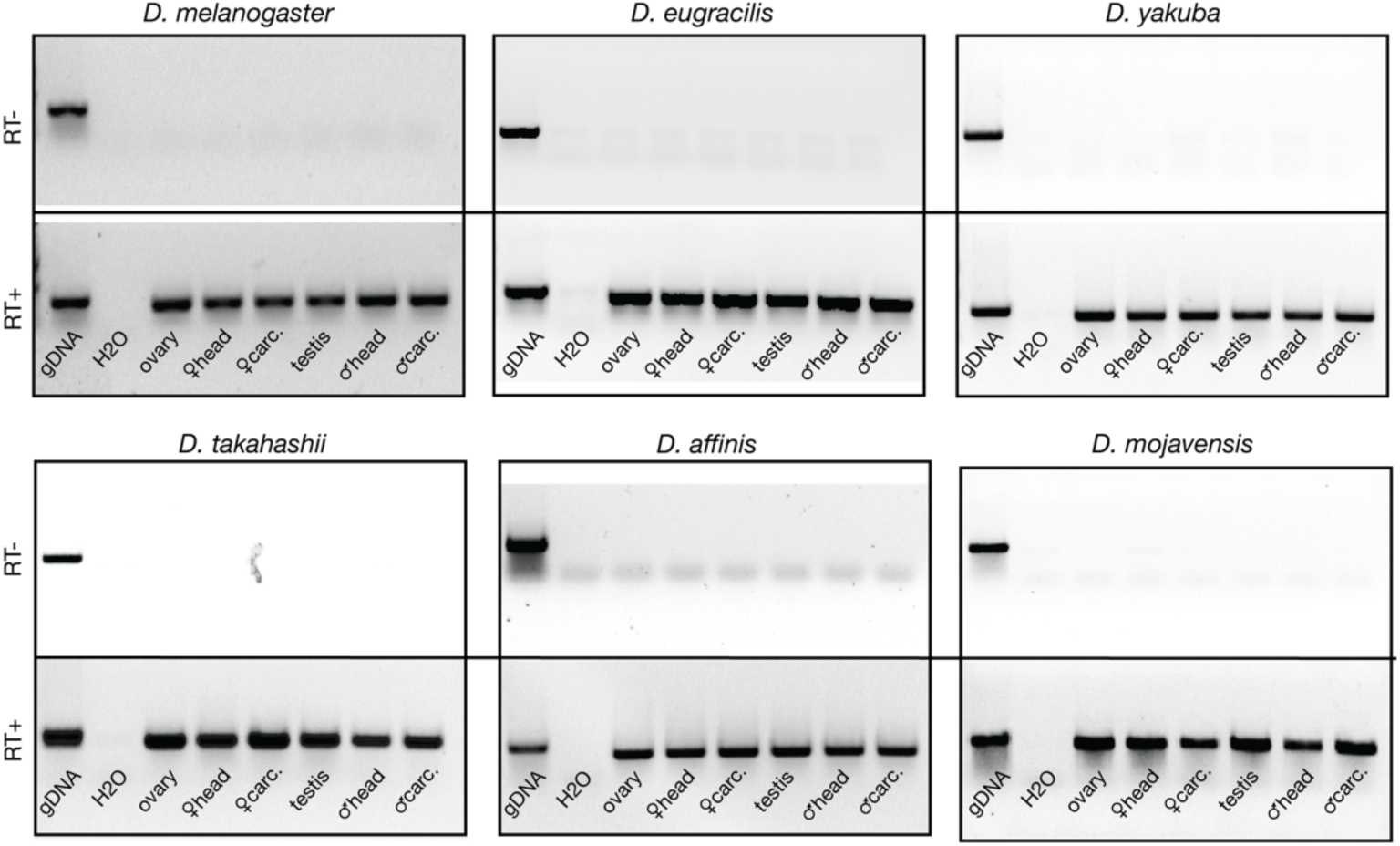
RNA templates for RT-PCR are free of genomic contamination. PCR products corresponding to the housekeeping gene, *rp49*, amplified across a panel of cDNA prepared with (bottom panels) or without (top panels) the reverse transcriptase enzyme to assess genomic contamination. gDNA = genomic DNA control, “carc.” = carcass.

**Figure S5.**
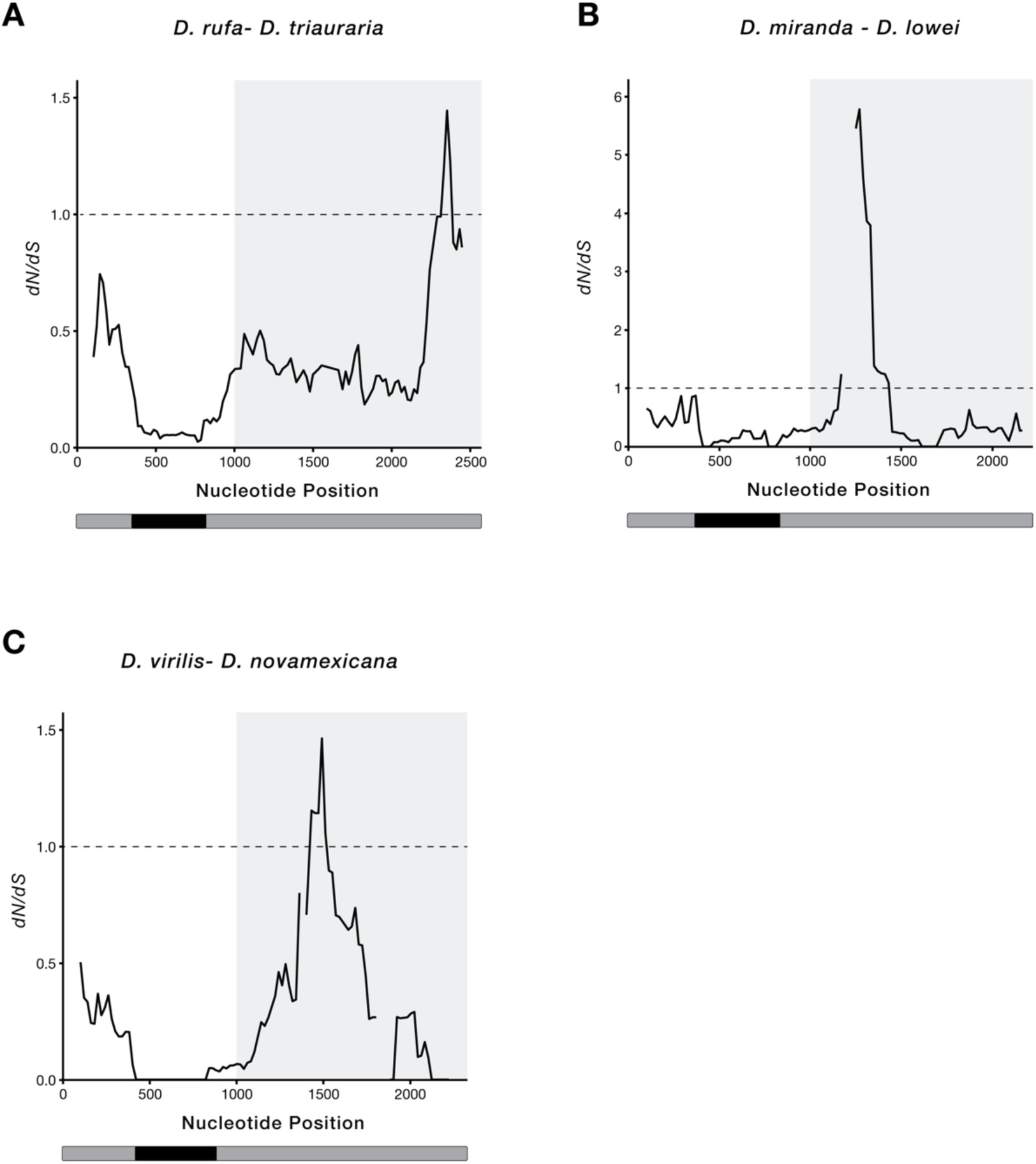
Peaks of *dN/dS* overlap the C-terminus of MH across multiple species pairs. Sliding window analysis of *dN/dS* across the *mh* coding region from (**A**) *D. rufa* and *D. triauraria* (*melanogaster* group), (**B**) *D. miranda* and *D. lowei* (*obscura* group), and (**C**) *D. virilis* and *D. novamexicana* (*virilis-repleta* group). Windows without diverged synonymous sites are represented as gaps. The conserved SprT domain is annotated in black. The C-terminal region, which includes the Spartan/MH regulatory domains, are denoted by a light gray box. Estimates of *dS* are <0.25 for all species pairs.

## SUPPLEMENTARY TABLES

**Table S1.** Results of the polarized McDonald-Kreitman Tests across *mh*, *CG2694*, *CG1132*.

**Table S2:** Species used for BLAST and tissue-specific expression analysis (in bold).

**Table S3:** SprT domain sequences used to build phylogenetic trees.

**Table S4:** Primers used in this study.

## Notes

### Competing Interest Statement

The authors have declared no competing interest.

