## Supplementary material for "Recurrent duplication and diversification of a vital DNA repair gene family across Drosophila": Table S1

**Table S1. Results of the polarized McDonald-Kreitman Tests across *mh*, *Gcna*, *CG2694*, *CG1132*.** Counts of polarized synonymous and non-synonymous polymorphic and fixed sites in the coding sequence of Spartan family genes are reported below. The coding sequences of *D. yakuba* *mh*, *Gcna*, *CG2694*, *CG1132* were used as an outgroup to polarize substitutions along the *D. melanogaster* and *D. simulans* lineages.

| Gene name |  |  | Polymorphisms | Fixations | p-value |
| --- | --- | --- | --- | --- | --- |
| *mh* | Synonymous | *D. melanogaster* lineage | 4 | 36 | *mel*-lineage: 0.72  *sim*-lineage: 0.41 |
|  |  | *D. simulans* lineage | 12 | 17 |  |
|  | Non-Synonymous | *D. melanogaster* lineage | 4 | 48 |  |
|  |  | *D. simulans* lineage | 8 | 20 |  |
| *CG2694* | Synonymous | *D. melanogaster* lineage | 13 | 23 | ***mel*-lineage: 8.9e-5**  ***sim*-lineage: 3.1e-6** |
|  |  | *D. simulans* lineage | 24 | 14 |  |
|  | Non-Synonymous | *D. melanogaster* lineage | 3 | 56 |  |
|  |  | *D. simulans* lineage | 18 | 70 |  |
| *CG11322* | Synonymous | *D. melanogaster* lineage | 14 | 22 | *mel*-lineage: 0.21  ***sim*-lineage: 0.003** |
|  |  | *D. simulans* lineage | 23 | 10 |  |
|  | Non-Synonymous | *D. melanogaster* lineage | 8 | 25 |  |
|  |  | *D. simulans* lineage | 18 | 33 |  |
