## Supplementary material for "Recurrent duplication and diversification of a vital DNA repair gene family across Drosophila": Table S4

| Forward | Reverse | Purpose |
| --- | --- | --- |
| CGTCAAACAGGTGACCATTC | GGTAAACGGTGATGTTGGTG | Primers used to amplify *mh* in *D. melanogaster* |
| GAGCCCAATTCTCAAGTTGC | TTGCGAAGTTTTCCCGATAG | Primers used to amplify *Gcna* in *D. melanogaster* |
| CCTGCTTCTGCACAGCATAG | GATTTCAAGTCGGCCAAGC | Primers used to amplify *CG2694* in *D. melanogaster* |
| GCGATCTTCCACTGTGGCCTTC | AACGCCATATTCGTGGAGTGCG | Primers used to amplify *CG11322* in *D. melanogaster* |
| CAGGATTCAGGCCCTCATAG | GCACTGGCTGTTTCTTTGG | Primers used to amplify *mh* in *D. yakuba* |
| GAAGAAGGCTCCCCTGAGAC | TCGTGTGAATGGCTTTTGAG | Primers used to amplify *Gcna* in *D. yakuba* |
| CGTGGATACCGACATGGAG | CGACCGCGATACTACAACAC | Primers used to amplify *CG2694* in *D. yakuba* |
| TCCAGGCCATTGTCTACTCC | GCGACTGAACGAATCCGTAG | Primers used to amplify *CG11322* in *D. yakuba* |
| GGCAATCGACATTTTAAGGAG | CTTAAAGTTGGGCCCATGAC | Primers used to amplify *mh* in *D. eugracilis* |
| ATGGAAGATGTGGGCTCAAC | CCGTATTTTCTCCACCTTGC | Primers used to amplify *Gcna* in *D. eugracilis* |
| CACTGCTGCACGAGATGTG | AATCCATCGTGCCGTAAGAG | Primers used to amplify *CG2694* in *D. eugracilis* |
| GAGGTGACCATACGGCTGAG | CTCCATGATGCGCTTGAAC | Primers used to amplify *mh* in *D. takahashii* |
| CTGCCACAACTACAGCATCG | TTCCCTGCTTGTCCTTCTTG | Primers used to amplify *Gcna* in *D. takahashii* |
| TCCTGTGGGAGAACATCAGC | TGTACTTGAAGGTGGCATCG | Primers used to amplify *CG2694* in *D. takahashii* |
| CCGGAGATGCTCATCAAAAT | CTCGAATTCAATCCGTCCAT | Primers used to amplify *CG2694 B*  in *D. takahashii* |
| GAGCCGCTGCTAAAGTTACG | GAAGGGATGACGGTTCTGAC | Primers used to amplify *mh* in *D. affinis* |
| CGGACGTGTTCAACAATGAG | GTAATGGTGGGCAAGTCTGG | Primers used to amplify *Gcna* in *D. affinis* |
| CTGGGTTTCAGCTCTTCAGG | TAGGAGCTACGGGCACAGAG | Primers used to amplify *CG2694* in *D. affinis* |
| TTCAGGCGAATTCCACCTAC | CCTTAGCCAGAGCCACTTTG | Primers used to amplify *CG2694 H* in *D. affinis* |
| ATCAGCGTGGTAACCGATTC | ATCCACCTCATCGTGAAAGG | Primers used to amplify *mh* in *D. mojavensis* |
| TCAGCCGTGAGTTCTGTGAC | ATCCGAGATGGTCTTGTTGC | Primers used to amplify *Gcna* in *D. mojavensis* |
| CTGCCGGAAATAGTGGATTG | TATAGACGTCGTCGGTGCAG | Primers used to amplify *CG2694* in *D. mojavensis* |
| TCGAAGCTGATGTTGAGCTG | GCCTGAGTGAAGGAAACGAC | Primers used to amplify *CG2694 M* in *D. mojavensis* |

**Table S4. Primers used in this study.**
